# Parallel and non-parallel features of adaptive radiation in Yucatán pupfishes

**DOI:** 10.1101/2025.11.17.688971

**Authors:** Matthew C. Kustra, David Tian, M. Fernanda Palominos, Feifei Guo, Dylan Chau, Oskar Golwala, HoWan Chan, Andrés Alvarez Zapata, Reyna Guadalupe Cetz Paredes, Frida Ximena Cortés Sánchez, Sonia Gabriela Hernández, Adan Fernando Mar-Silva, Fernando Mex, Charles Tralka, Maribel Badillo-Alemán, Juan J. Schmitter-Soto, Carlos A. Gracida-Juárez, Christopher M. Martinez, Jairo Arroyave, Christopher H. Martin

## Abstract

Understanding the extent of parallelism across adaptive radiations remains a central problem in evolutionary biology. We used whole-genome resequencing of 123 individuals to compare the adaptive radiation of *Cyprinodon* pupfishes in Lake Chichancanab, Mexico, to an independent radiation of San Salvador Island (SSI) pupfishes in the Bahamas, and assess the repeatability of adaptive genetic architecture, sources of adaptive variation, and stages of selection. Despite rapid craniofacial divergence of trophic specialists within 8-15 kya, only two candidate genes (0.5%; 2/426) were shared between Caribbean radiations. Although adaptive introgression played a major role in SSI, we found minimal evidence of adaptive introgression in Chichancanab, likely due to the geographic isolation of this inland lake. Instead, de novo mutations provided a substantial source of adaptive variation (30.6%) for the endemic zooplanktivore, 15 times higher than the endemic scale-eater on SSI. However, in parallel with SSI, we found strong evidence that adaptive divergence occurred in stages, first on regulatory and standing genetic variation, then on de novo and nonsynonymous mutations. Consistent with adaptive variants near opsin and spermatogenesis genes, functional categories unique to Chichancanab, we found greater visual acuity and divergent sperm morphology in lab-reared zooplanktivores relative to generalists using laboratory assays. Consistent with extensive adaptive de novo mutations in *WNT10A* and rapid diversification of tooth size in the zooplanktivore, we found that experimental inhibition of the Wnt pathway in generalists resulted in narrower oral teeth. We conclude that de novo mutations, not introgression, can drive rapid adaptive radiations in isolated environments.

## Introduction

Much of biodiversity results from adaptive radiation—when a clade diversifies rapidly into three or more new species (1–5). Young adaptive radiations provide excellent systems for investigating speciation and adaptation (3, 6–9). However, it remains unclear to what degree radiations occur in parallel given similar starting positions on the adaptive landscape with similar levels of genetic diversity, gene flow, and ecological opportunity (10, 11).

Beneficial de novo mutations can provide novel variation to reach new fitness peaks on the adaptive landscape during adaptive radiation (12, 13). However, the longer time needed for de novo beneficial mutations to arise contradicts the short timescales of rapid divergence (11, 14, 15). Hybridization can introduce large amounts of genetic variation and novel phenotypes (transgressive segregation) through adaptive introgression and hybrid swarms, enabling populations to reach previously inaccessible fitness peaks (16). Support for an important role of hybridization in adaptive radiations has been demonstrated both theoretically (17, 18) and empirically in many systems (3, 19–27). However, we still do not know the relative importance of different sources of genetic variation for triggering and sustaining adaptive radiations across time and space.

Caribbean pupfishes are a remarkable system for assessing the repeatability of adaptive radiations due to two independent, recent adaptive radiations, each containing a generalist and multiple trophic specialists within isolated lake environments that have historically lacked predatory fishes and most competitors except *Gambusia*. The San Salvador Island (SSI), Bahamas radiation proceeded predominantly via the reassembly of pre-existing standing genetic variation found throughout the Caribbean, with substantial contributions of adaptive introgression to both the molluscivore (durophage) and scale-eating (lepidophage) specialists. Only 0% and 2% of adaptive mutations in the molluscivore and scale-eater, respectively, were de novo mutations unique to SSI (26, 28–32). However, it was previously unknown how a second radiation in Lake Chichancanab, Mexico, drawing from the same Caribbean pool of standing genetic variation, unfolded over the same timeframe.

Here, we use whole-genome resequencing of 123 individuals to provide insight into the genomic architecture and genetic origins of the Lake Chichancanab *Cyprinodon* pupfish radiation (33). This recent adaptive radiation (∼8000 years old (34)) consisted of at least five species (or up to seven (35)) historically (33, 35, 36), including a detritivore (*C. beltrani*), piscivore (*C. maya*), rocky substrate specialist (*C. verecundus*), bivalve specialist (*C. labiosus*), and zooplanktivore (*C. simus*). At the time of description (33), the relative abundances of these endemic species were 68-85% *C. beltrani*, 6-18% *C. maya*, 2-13% *C. labiosus*, and <1% *C. simus* and *C. verecundus*. However, due to recent colonization by invasive African tilapia (*Oreochromis niloticus x mossambicus*) (37, 38) and by several species native to surrounding basins but not Chichancanab (*Astyanax angustifrons, Mayaheros urophthalmus, Poecilia mexicana*; pers. obs.), all trophic specialist species now appear to be extinct since our collections in 2022, with only the generalist, *C. beltrani*, remaining in 2024. Despite exhibiting one of the fastest rates of trait diversification of any vertebrate in tooth size (130 times faster than background rates across *Cyprinodon* (29)) and rapid divergence in other craniofacial morphology (Figure 1A; S1), previous genetic studies of microsatellite data and mtDNA suggest that this radiation has low genetic differentiation (39, 40), enabling the identification of highly differentiated loci among these species.

**Figure 1.**
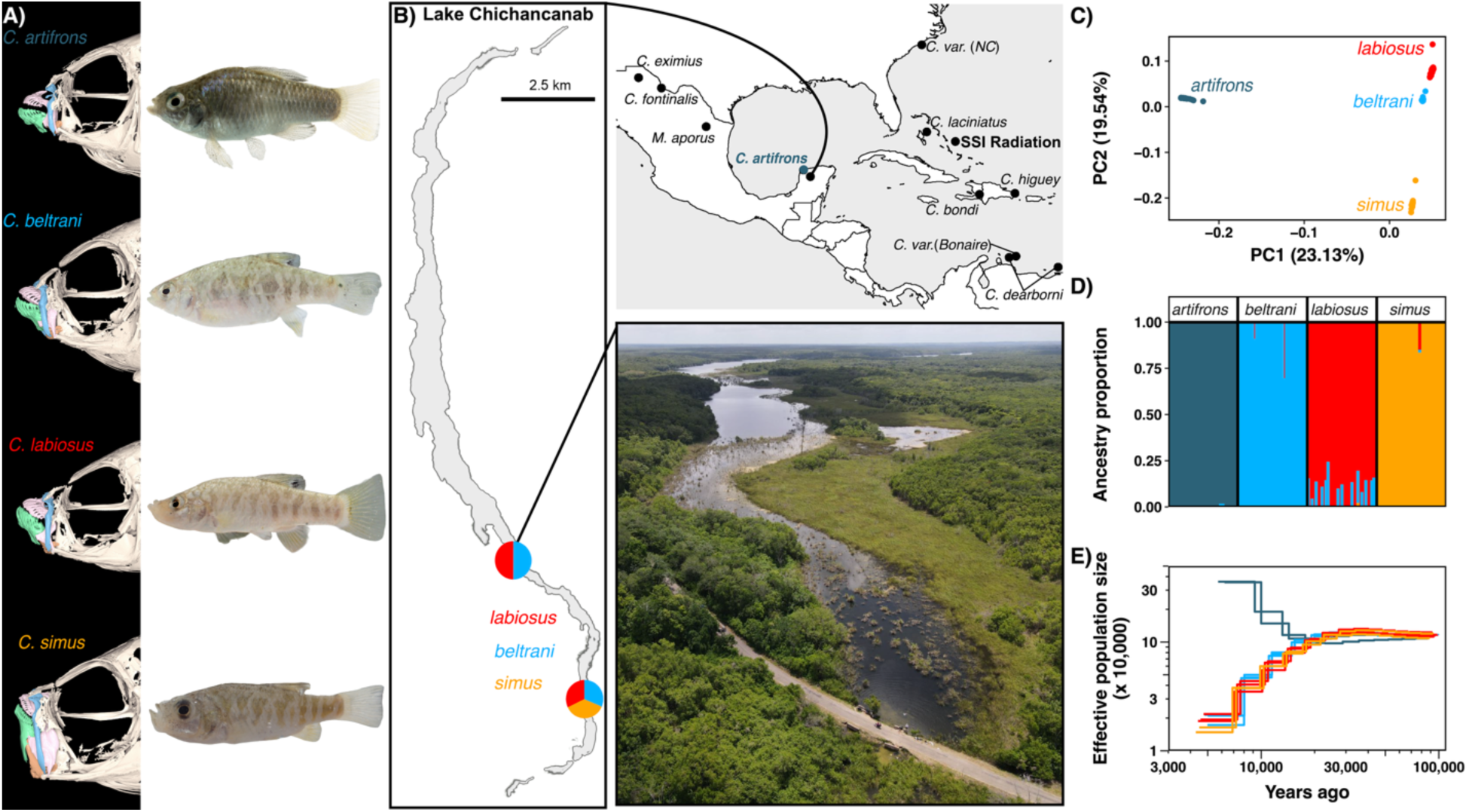
Yucatán *Cyprinodon* pupfishes show strong genetic differentiation. (A) µCT scans of Yucatán pupfish with maxilla colored in blue, premaxilla in pink, dentary in green, and articular in orange (see Figure S1 for different views). (B) Map of the Caribbean showing the sampling sites for all species in our dataset with an inset map of Lake Chichancanab and an aerial view image from one of the sampling sites. (C) Principal component analysis using a set of linkage disequilibrium pruned SNPs. (D) Estimated ancestry proportions for each individual inferred using *ADMIXTURE* with K = 4 (see Figure S3 for K=2-3) from the linkage-pruned dataset. (E) Estimated demographic history with *MSMC2* on a log scale. Each colored line represents an individual, with colors corresponding to different species.

## Results

### Genetic differentiation of the species flock

To measure population structure of the Yucatán *Cyprinodon* pupfishes, we conducted whole genome resequencing of the species flock and the coastal sister species (*n* = 15 *C. artifrons*, *n* = 35 *C. beltrani*, *n* = 35 *C. labiosus*, and *n* = 17 *C. simus*) using specimens collected from the wild in 2022 (Figure 1B). We aligned these samples, as well as eleven other previously sequenced Caribbean pupfish species, to the high-quality *Cyprindon nevadensis mionectes* reference genome (GCA_030533455.1). After calling and filtering variants, our dataset included 23,960,536 SNPs (12.4x mean depth). Using a linkage disequilibrium pruned set of SNPs (*n* = 5,476,855 SNPs), we used principal component analysis and *ADMIXTURE* to visualize population structure within Yucatán pupfishes. We found that all species formed distinct, non-overlapping genetic clusters, with *C. artifrons* being the most genetically divergent from and sister to the Chichancanab species (Figure 1C). Within the Chichancanab species flock, *C. simus* was more genetically divergent from *C. labiosus* and *C. beltrani* (Figure 1C). These results were also supported by genome-wide *F_ST_* estimates and a phylogenetic tree constructed with *ADMIXTOOLS2* (Table S1; Figure S2). Four clusters corresponded to the four different species, as determined by *ADMIXTURE* (Figure 1D). All *C. artifrons* individuals showed minimal evidence of admixture with Chichancanab species (Figure 1D; Figure S3).

### Demographic history

Using *MSMC2* for Chichancanab species with a mutation rate of 1.56 x 10^-8^ substitutions per base pair (estimated from a Caribbean *Cyprinodon* species (26)), we found that the three Chichancanab species (*C. beltrani*, *C. labiosus*, and *C. simus*) diverged from the coastal sister species *C. artifrons* at the end of the Pleistocene 10-15 kya, consistent with receding sea levels resulting in the formation of Lake Chichancanab (34). After divergence from *C. artifrons*, the Chichancanab species flock showed a similar demographic history, characterized by a sharp decline in effective population sizes, consistent with endemic sympatric species within an isolated lake (Figure 1E).

### Adaptive loci within trophic specialists

Although genome-wide differentiation among Chichancanab species was moderately low (*F_ST_* ranging from 0.032 to 0.09; Figure 2A; Table S1), we were able to identify highly differentiated regions of the genome within the trophic specialist zooplanktivore *C. simus*. We identified 19 fixed and 1,127 nearly-fixed (*F_ST_* >0.95) SNPs in *C. simus* relative to other Chichancanab pupfishes (Tables S2-S3). In contrast, we found zero fixed or nearly-fixed SNPs within the generalist *C. beltrani* or bivalve-specialist C. *labiosus*. However, we were able to identify some moderately differentiated SNPs within *C. labiosus* (*n* = 97) and *C. beltrani* (*n* = 13) relative to the other Chichancanab species at an *F_ST_* threshold of 0.8 (Table S2).

**Figure 2.**
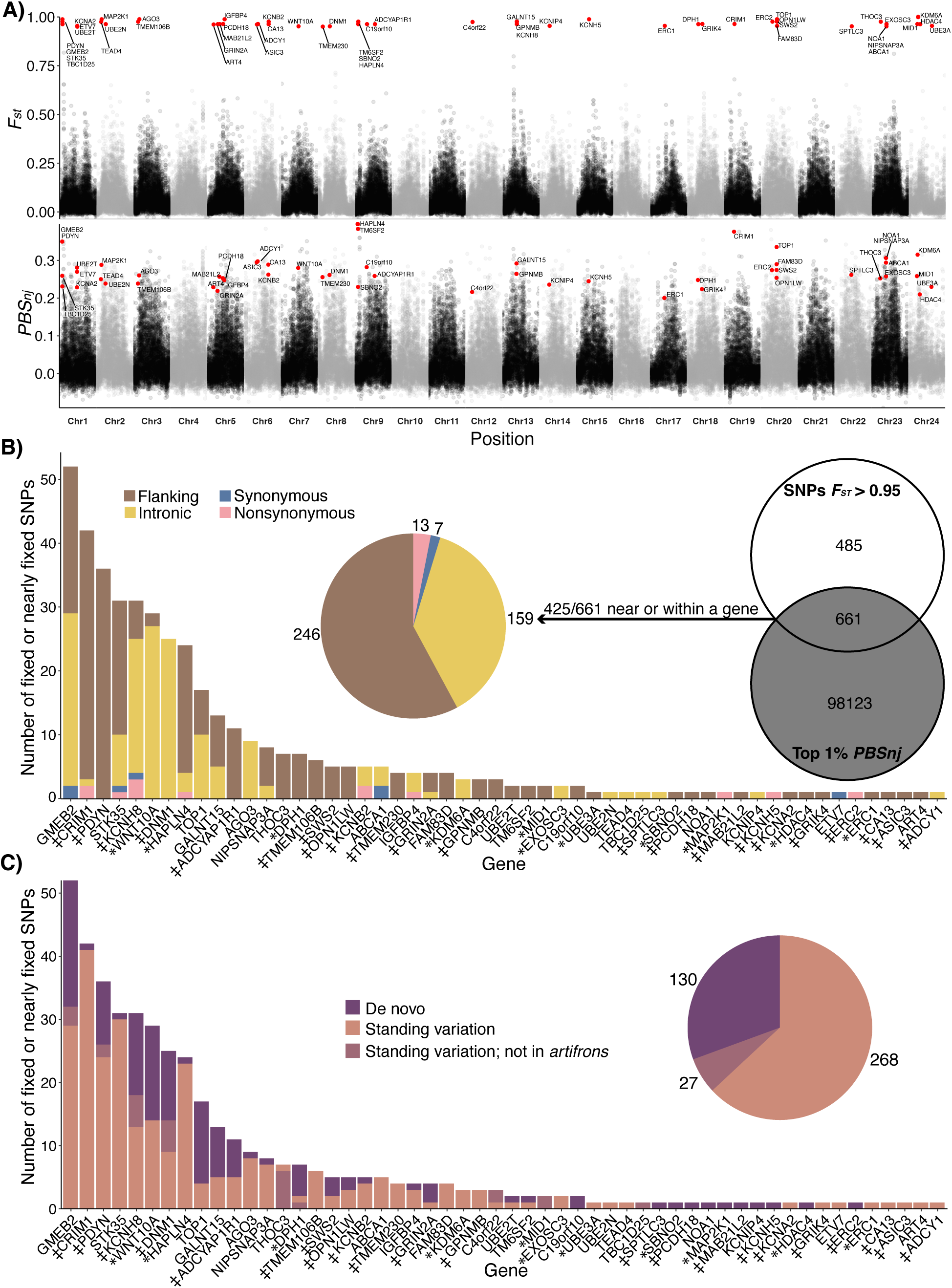
Architecture and source of adaptive genetic variation in the zooplanktivore. (A, top row) *F_st_* between *C. simus* and *C. beltrani* + *C. labiosus* calculated in non-overlapping 10 kb windows. *F_st_* values for candidate SNPs (red circles) were calculated separately per locus to show the location of fixed variants. (A, lower row) Population branch statistics (*PBS_nj_*) with *C. simus* as the target population versus *C. beltrani*, *C. labiosus*, and *C. artifrons* in 10-kb non-overlapping windows. Candidate genes within 20 kb of fixed or nearly-fixed (Fst > 0.95) SNPs and within *PBS_nj_* 1% outlier windows are highlighted in red and labeled. Alternating colors represent different chromosomes. (B) Candidate adaptive variants near or within a gene were defined as SNPs that were fixed (*n* = 17) or nearly fixed (*n* = 1,129; *F_st_* > 0.95) and occurred within a 1% population branch statistic (*PBS_nj_*) outlier window for *C. simus* (Venn diagram). Pie chart and bar graphs indicate the proportion of these 425 adaptive variants that were found within the flanking, intronic, or coding regions. (C) Pie chart and bar graphs indicate the proportion of these 425 variants that were detected only within Chichancanab (de novo: purple), as standing genetic variation in coastal sister species *C. artifrons* (pink), or in other Caribbean species, but not *C. artifrons*. * Indicates genes with a craniofacial annotation; ‡ indicates genes with neural or sensory annotation.

To search for signals of selection, we calculated the normalized four-taxon population branch statistic (*PBS_nj_*) across 10-kb non-overlapping windows because linkage disequilibrium decayed at this distance (Figure S4). Simulation studies demonstrate that normalized *PBS* is effective for identifying both hard and soft selective sweeps (41). Out of the 1,146 SNPs fixed or nearly fixed within *C. simus*, 661 were within a strongly supported selective sweep window (top 1% *PBS_nj_* outlier (41)), including 425 SNPs within 20 kb of the first or last exon of an annotated gene. These 425 candidate adaptive SNPs (“adaptive variants”) were located in the proximity of, or within, 55 genes (Figure 2A), distributed across 19 out of 24 chromosomes (Figure 2A).

Adaptive variants associated with the same gene were often distributed across multiple introns or exons and spaced apart by 1kb or more (Figure S5, Table S4).

### Source and function of adaptive genetic variation in the zooplanktivore

We next characterized the location of the 425 candidate adaptive variants near or within a gene in the zooplanktivore *C. simus*. More than 95% were in intronic or flanking regions (20-kb upstream or downstream of a gene; Figure 2B). Within coding regions, 13 out of 20 adaptive variants were nonsynonymous mutations (Figure 2B). The number of adaptive variants per gene followed an exponential distribution, with a handful of genes containing many variants and most genes with only a single adaptive variant (Figure 2B). We functionally characterized these genes by manually checking online databases and doing a literature search (Table S3; Figure 2B). 32 out of the 55 candidate genes were associated with craniofacial development (*n =*11; 20%) or sensory/neural function (*n =* 21; 38.2%). We manually checked if any of these genes overlapped with the 426 unique candidate genes identified in the SSI radiation (26, 28, 30, 32, 42–46), and we found only two overlapping candidate genes (*DNM1* and *SPTLC3*), both of which are associated with motor neuron development (47, 48) and are scale-eater-specific candidate genes.

Next, we characterized the source of the variants and found that 63% of the adaptive SNPs in *C. simus* also existed as standing genetic variation in the coastal Yucatan population of *C. artifrons* (“standing variation”), 31% were *de novo* variants found only in Chichancanab, and 6% were detected as standing genetic variation in other pupfish species but not *C. artifrons* (Figure 2C).

### Limited adaptive introgression within Chichancanab pupfishes

To test for introgression into the Chichancanab radiation, we used *f*-statistics calculated in *Dsuite* (49) to examine gene flow from 11 additional *Cyprinodon* species from across the clade and the sister species to the genus *Megupsilon aporus*. Within Yucatán pupfishes, we found evidence of introgression between *C. artifrons* and both *C. beltrani* and *C. labiosus*, with the strongest signal in *C. beltrani* (Figure 3A, B). We also found evidence for introgression between *C. beltrani* and most outgroup species examined, including the sister species to *Cyprinodon*, *Megupsilon aporus*. Relative to *C. beltrani*, *C. labiosus* and *C. simus* also showed evidence of introgression with pupfishes endemic to the Chihuahua desert (*C. eximius* and *C. fontinalis*; Figure 3A, B). Furthermore, we also found evidence of introgression between the two sympatric radiations (*C. variegatus* from SSI) and *C. simus* (Figure 3A, B). However, additional *f-*branch statistics indicate that all patterns of excess allele sharing between these outgroup species besides *C. artifrons* and the Chichancanab species flock likely resulted from ancient introgression events before the colonization of Lake Chichancanab (Figure 3C; S6).

**Figure 3.**
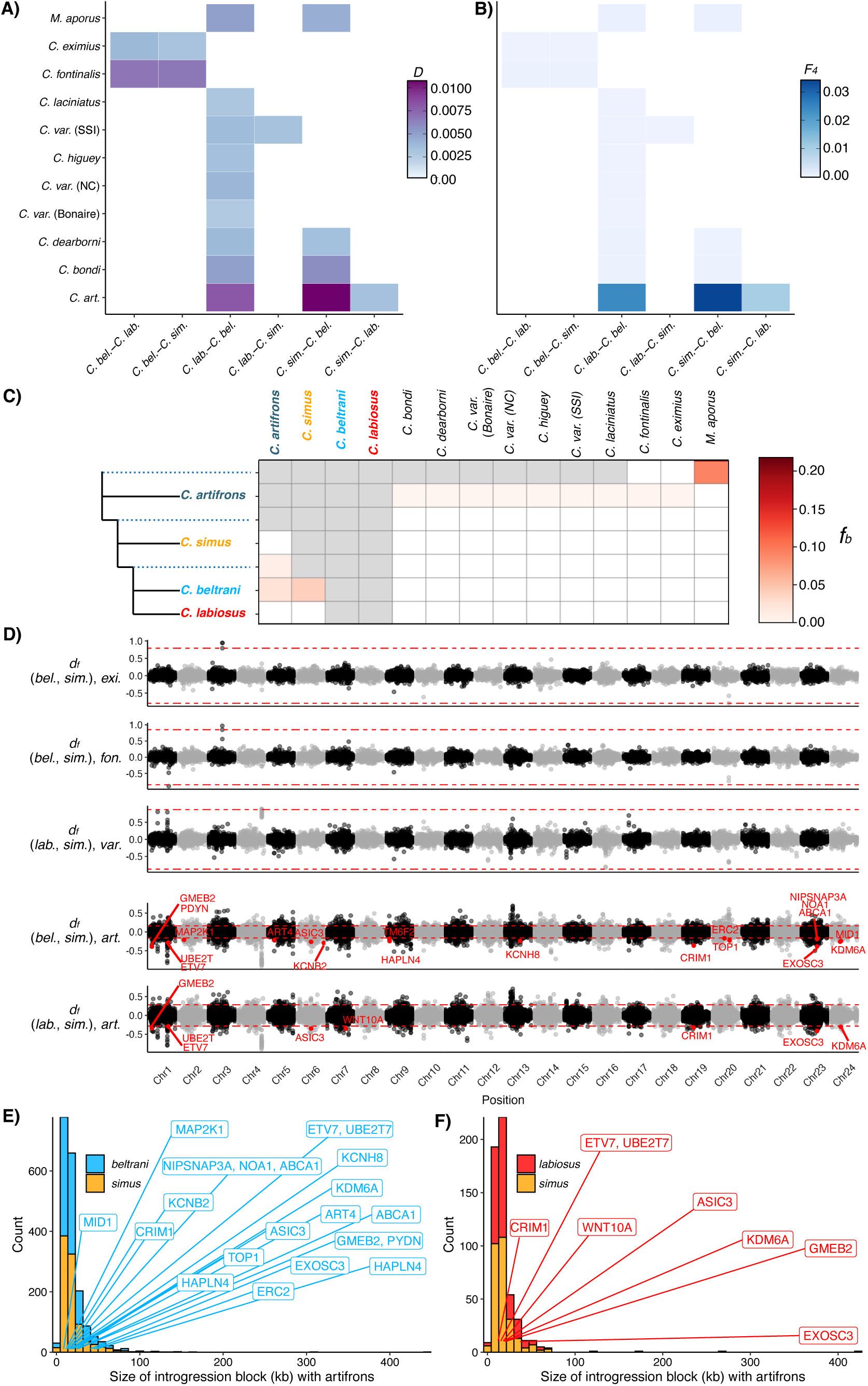
Minimal evidence of adaptive introgression into Chichancanab pupfishes. Heatmaps of (A) Patterson’s *D*-statistics and (B) *F_4_* ratios for P1-P2 species combinations within the Chichancanab radiation along the x-axis and P3 focal species along the y-axis. All four-taxon trees used *Cualac tessellatus* as the outgroup (P4). Only significant (*Z*-score > 3) results are colored; combinations not relevant to the Chichancanab radiation are not shown. (C) *f-*branch (*f_b_*) statistics indicate introgression occurring before the Chichancanab radiation. Red boxes indicate significant introgression between P3 species (top row) and the focal species or internal branch (dashed lines) on the left. Non-significant *f_b_* statistics are indicated by white boxes. Grey boxes represent comparisons that cannot be made since *f_b_* cannot be calculated for introgression between sister taxa. See Figure S6 for full *f-*branch graphs. (D) Genome-wide plot of the distance fraction (*d_f,_*, an introgression test statistic (50)) calculated in non-overlapping windows of 91 SNPs (approximately 10 kb) for species trios that showed evidence of introgression with *C. simus. d_f_* outlier windows that contained a candidate gene under selection in *C. simus* are colored red and labeled with the gene name. The y-axis label gives the introgression topology: (P1, P2), P3. Positive values indicate regions of introgression between *C. simus* and P3 (from top to bottom: *C. eximius*, *C. fontinalis*, *C. variegatus* (SSI), and *C. artifrons* in the bottom two subplots); negative values indicate regions of introgression between P1 (*C. beltrani* or *C. labiosus*) and P3. Red dashed lines indicate outlier cutoffs, which were determined empirically by the overall percentage of genome-wide introgression estimated from the *F_4_* ratios in panel B. (E, F) Distribution of the size of *d_f_* outlier introgression tract lengths (in 10 kb windows) between *C. artifrons* and Chichancanab species. The introgression tracts containing candidate adaptive genes in *C. simus* are labeled with gene names. SSI: San Salvador Island, Bahamas; NC: North Carolina, USA.

To test for adaptive introgression into *C. simus*, we next examined whether the set of adaptive variants (Figure 2) occurred within outlier introgression windows using sliding window scans of distance fraction (*d_f_* (50)) for species that had significant genome-wide introgression into *C. simus* (Figure 3A, B). We found that none of the outlier introgression windows overlapped with *C. simus* adaptive variants (Figure 3D).

Next, we performed *d_f_* scans for introgression between *C. artifrons* and Chichancanab species. Although there were outlier introgression regions between *C. simus* and *C. artifrons* (positive values *d_f_*), these regions did not overlap with *C. simus* adaptive variants (Figure 3D). However, the position of several of these *C. simus* adaptive variants overlapped with outlier introgression windows for both *C. labiosus* and *C. beltrani* but not *C. simus* (Figure 3D). The number of outlier introgression windows was much lower in *C. simus* relative to *C. labiosus* and *C. beltrani* (Figure 3E, F). Furthermore, both *C. labiosus* and *C. beltrani* populations contained several introgression tracts that were much longer than *C. simus* tract lengths, indicating much more recent introgression events in these species (Figure 3E, F).

### The stages of adaptive divergence of the zooplanktivore

We used *starTMRCA* (51) to estimate the timing of selective sweeps for the set of candidate adaptive variants near genes within the zooplanktivore *C. simus*. We estimated that most selective sweeps occurred in a staggered sequence and not simultaneously after colonization of Lake Chichancanab (Figure 4A)(34). There were also a few older sweeps that we estimate may predate the age of Lake Chichancanab and divergence from the coastal sister species *C. artifrons* (Figure 1B). Overall, we observed no clear temporal stages of adaptive phenotypic divergence related to craniofacial genes (asterisks) or neural/sensory genes (‡; Figure 4A). Instead, we found that selection on these two broad functional categories occurred throughout the period of divergence of *C. simus* following the colonization of Lake Chichancanab, contrary to behavior-first, then craniofacial stages of adaptation (52) found in the SSI radiation (26).

**Figure 4.**
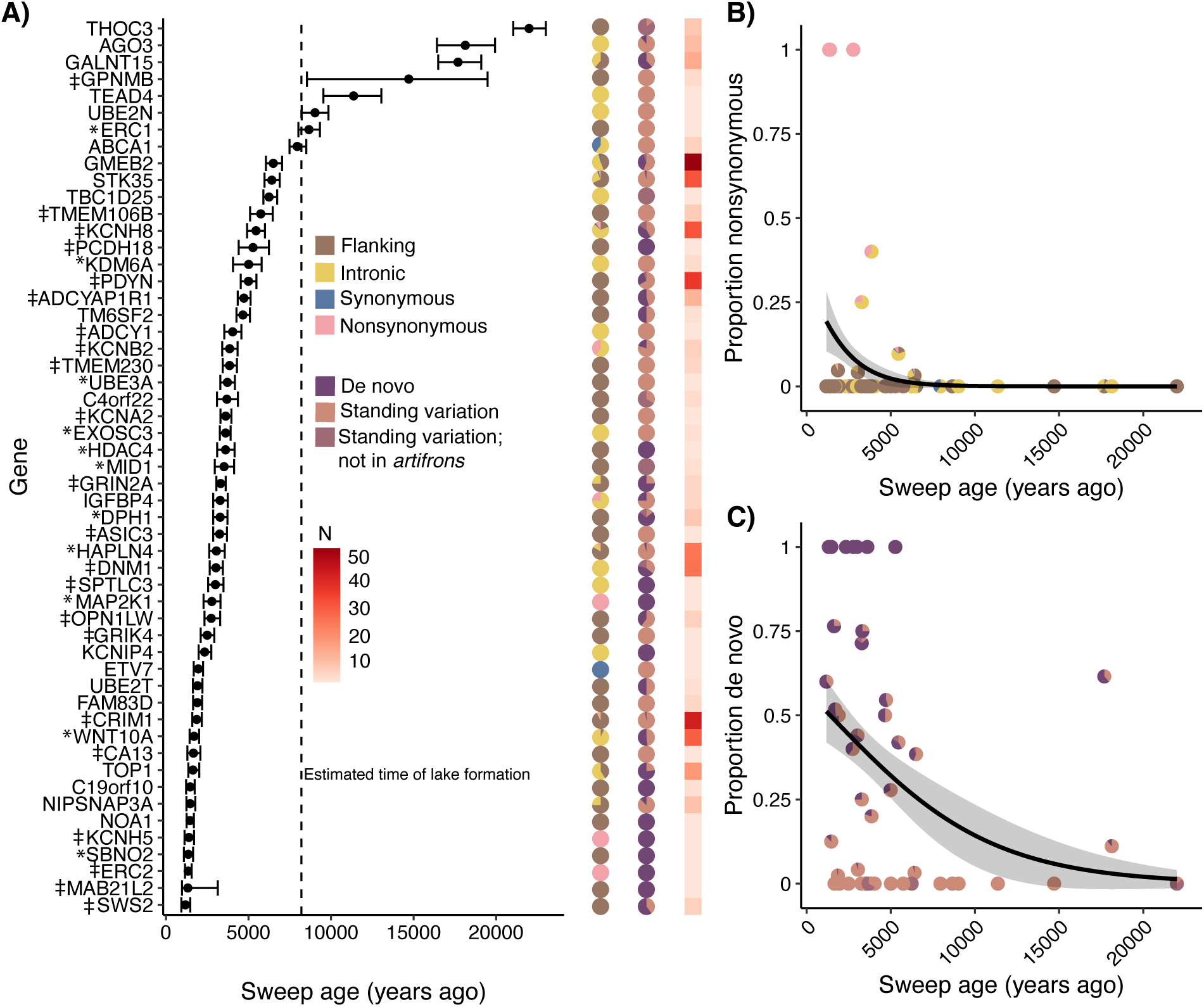
Stages of adaptive divergence in the zooplanktivore. (A) Points indicate median sweep age with 95% credible intervals. The dashed line indicates the approximate age of Lake Chichancanab (34). For each sweep, the first column of pie charts shows the proportions of flanking/intronic/synonymous/nonsynonymous, second column of pie charts shows the proportions of de novo/standing variation, and third column of squares shows the number of candidate adaptive variants per gene (from Figure 2). For genes with multiple adaptive variants, we used the variant with the median position as the focal position. * Indicates genes with a craniofacial annotation; ‡ indicates genes with neural or sensory annotation. (B) The proportion of nonsynonymous adaptive variants is significantly larger in more recent selective sweeps, indicated by the best-fit quasibinomial regression line; each pie chart represents a single candidate gene. (C) The proportion of de novo variants is significantly larger in more recent selective sweeps; each pie chart represents a single candidate gene.

Next, we used generalized linear regressions to test if sweep age was correlated with the proportion of nonsynonymous adaptive variants, de novo adaptive variants, or the overall number of adaptive variants per gene region. We found that both nonsynonymous adaptive variants (χ^2^ = 5.852, *p* = 0.016) and de novo mutations (χ^2^ = 6.221, *p* = 0.013) were significantly more likely to occur in more recent selective sweeps during the final stage of adaptive divergence in *C. simus* (Figure 4B, C). However, nonsynonymous mutations were not significantly more likely to be de novo mutations found only in Chichancanab species (All adaptive variants: χ^2^ = 0.102, *p* = 0.749; singleton adaptive variants: χ^2^ = 2.010, *p* = 0.156), and there was no correlation between estimated sweep age and the number of candidate adaptive variants per gene region (χ^2^ = 0.111, *p* = 0.740).

### Increased visual acuity in the zooplanktivore is consistent with vision-related candidate adaptive genes

We identified several *C. simus* adaptive variants associated with eye development (*ASIC3*(53), *CRIM1*(54), *MAB21L2*(55)) and opsin (*OPN1LW*, *SWS2*) genes. Because the trophic specialization of *C. simus* requires visually locating zooplankton in the water column before precise suction-feeding strikes, we hypothesized that this species would have higher visual acuity than the detritivore, *C. beltrani*. We conducted optomotor response trials on lab-reared individuals from both species (F1 *C. beltrani* and long-term laboratory colony of F10+ *C. simus*) and found that *C. simus* has a much stronger optomotor response than *C. beltrani* in response to black-and-white vertical bars spinning at a constant rate of approximately 90 rotations per minute (Figure 5A, B; Wilcoxon signed-rank test: *W* =1, *p* = 0.009). There was no significant difference in rotations per minute between the species for the same individuals during control observation periods with no spinning (Figure S7A; *W*=14, *p* = 0.3105) or during positive control observation periods with an all-black background spinning at the same frequency (Figure S7B; *W* =9, *p* = 0.147).

**Figure 5.**
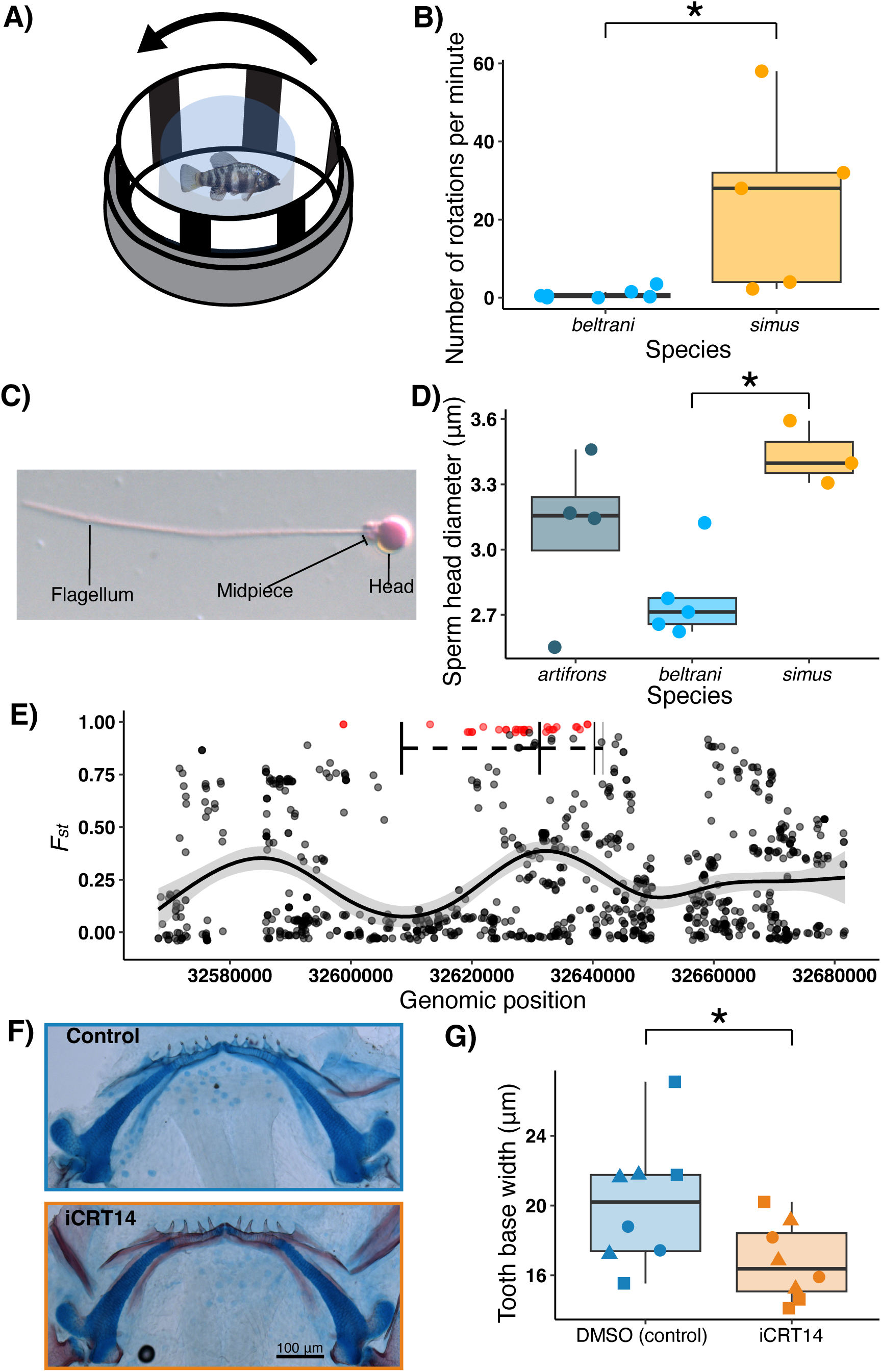
Divergence in visual acuity, sperm morphology, and tooth development reflects candidate adaptive genes in the zooplanktivore. (A) Diagram of the optomotor response experimental apparatus (adapted from (56)). (B) Boxplot and jittered points of the number of complete rotations fish swam per minute, indicative of their optomotor response to black-and-white vertical bars spinning at a constant rate. (C) Representative labeled image of a single *C. beltrani* sperm at 100x stained with rose bengal. (D) Boxplots with jittered points of sperm head diameter for *C. artifrons, beltrani,* and *simus.* Significant Tukey’s post hoc differences are shown with an asterisk. (E) Manhattan plot showing per-site *F_st_* of SNPs within 50 kb of *WNT10A*. Black rectangles indicate exons of *WNT10A*, and dashed lines indicate introns. Red points represent nearly fixed SNPs (*F_st_* > 0.95), and black points represent all other SNPs. The black line is the smoothed average *F_st_* in this region, with grey shading being the 95% confidence intervals. (F) Representative images of *C. beltrani* larvae that were treated with DMSO (control; top) or with iCRT14 Wnt inhibitor + DMSO (bottom) and stained with alizarin red (bone) and alcian blue (cartilage). (G) Boxplot and jittered points of average tooth base width of the left and right teeth closest to the mandibular symphysis for *C. beltrani* larvae treated with a DMSO control or Wnt inhibitor (iCRT14) + DMSO. Shapes represent three independent split-brood replicates.

### Divergent sperm morphology of the zooplanktivore is consistent with spermatogenesis-related candidate adaptive genes

Four candidate genes (*C4orf22*, *STK35*, *ERC2*, and *TBC1D25*) are known to affect spermatogenesis/fertility (57–60). We therefore examined whether three Yucatan species, raised in a common garden laboratory environment (F1 *C. artifrons,* F1 *beltrani,* and a long-term laboratory colony of F10+ C. *simus*), differed in sperm morphology using differential interference contrast microscopy. We found that species significantly differed in sperm head length (Figure 5D; *F* = 5.724, *p* = 0.025), but not sperm midpiece length (Figure S8A; *F* = 2.403, *p* = 0.147) or flagellum length (Figure S8B; *F* = 0.414, *p* = 0.673). *C. simus* had significantly larger sperm heads than *C. beltrani* using Tukey’s post hoc pairwise comparisons (*q* = 3.367; *p* = 0.0207; Fig. 5D).

### Inhibition of WNT10A alters tooth morphology in Chichancanab pupfish

We identified a large number of highly differentiated SNPs surrounding the well-characterized *WNT10A*, which controls tooth, hair, and skin development within the Wnt signaling pathway (Figure 2; Figure 5E (61–63)). This is consistent with the most exceptional morphological difference between *C. simus* and *C. beltrani,* extreme divergence in tooth size (130 times faster than background rates (29)).

To test for the importance of *WNT10A* in tooth development within pupfishes, we used iCRT14 to inhibit the Wnt / β catenin pathway downstream of WNT10A during metamorphosis in *C. beltrani* lab-reared fry (64, 65). Chemical inhibition from 8 dpf to 22 dpf resulted in narrower teeth than control fry treated with DMSO (χ^2^ = 4.840, *p* = 0.0278; Figure 5F, G). There were no significant differences in mean tooth length (χ^2^ = 0.455, *p* = 0.500; Figure S9) or number of ossified teeth when stained with alizarin red (χ^2^ = 0.114, *p* = 0.7357; Figure S10).

## Discussion

Here, we investigated the genetic architecture, sources of adaptive variation, and temporal dynamics of selection underlying speciation and trophic specialization in a young adaptive radiation of *Cyprinodon* pupfishes endemic to Lake Chichancanab, Mexico. By comparing these patterns to our previous work on a recent radiation of *Cyprinodon* pupfishes in SSI, Bahamas (26, 28–32), we discovered parallel and non-parallel features of these independent radiations in similar environments. Despite replicate radiations of Caribbean pupfishes in large, isolated, predator-free, saline lakes with few competitors, we found little parallelism between the radiations. Although craniofacial morphology diverged rapidly within trophic specialists in both radiations, we found almost zero overlap in the adaptive genes underlying trophic specialization (0.5%), potentially reflecting the divergent specialists within each radiation. In contrast to SSI, there was no behavior-first stage of adaptive divergence in Chichancanab specialists and, with the exception of craniofacial development, candidate adaptive gene ontologies potentially relevant to speciation were not shared between radiations, such as pigmentation genes in SSI versus opsin and spermatogenesis genes in Chichancanab. Unlike the SSI radiation and most other well-studied adaptive radiations (e.g., Malawi cichlids (20), Galápagos finches (66)), adaptive introgression likely played a minimal role in the Chichancanab radiation. We found no overlap between adaptive variants in the specialists and tracts of introgression. Instead, over 30% of the candidate adaptive variants in the zooplanktivore, *C. simus*, likely arose from de novo mutations compared to only 2% in the SSI scale-eater specialist and none in the molluscivore specialist. One outstanding parallel feature was a clear stage of refinement in the adaptive divergence of the zooplanktivore, marked by increased rates of de novo and nonsynonymous adaptive mutations occurring among the most recent selective sweeps in this species, similar to the scale-eating specialist on SSI. In contrast to *C. simus*, we identified no sufficiently differentiated SNPs within *C. labiosus*, the bivalve specialist, to identify candidate genes. Nonetheless, minimal divergence of this specialist in Chichancanab parallels the minimal divergence of the molluscivore on SSI, each in contrast to a more divergent trophic specialist.

### Adaptive introgression did not contribute to speciation of Chichancanab pupfishes

Genomic analyses have revealed that hybridization and adaptive introgression played a major role in many adaptive radiations (3, 11, 16, 19–27, 67, 68), including the *Cyprinodon* SSI radiation (26, 31). However, in Chichancanab, we found minimal evidence of introgression overall and did not detect any signatures of adaptive introgression following the colonization of Lake Chichancanab. Although our analyses indicate extensive introgression among *Cyprinodon* and *Megupsilon* lineages, we did not find any introgression events into the root of the radiation, nor has there been secondary gene flow from outgroup species other than the sister species to the radiation, *C. artifrons*. Secondary gene flow from *C. artifrons* was almost entirely detected with *C. beltrani* and, to a lesser extent, *C. labiosus.* Gene flow may be higher with these species because *C. artifrons,* a coastal benthic feeding detritivore, is less ecologically divergent from both *C. beltrani,* a benthic feeding detritivore, and *C. labiosus*, a benthic feeding bivalve specialist. Secondary gene flow between *C. artifrons* and both *C. beltrani* and *C. labiosus* overlapped with many genes under selection in *C. simus*, suggesting that secondary gene flow is helping increase similarities between the benthic detritivores within the lake and the coastal *C. artifrons* population rather than by bringing in adaptive variation for trophic specialization in *C. simus*.

These results are remarkably different from the *Cyprinodon* adaptive radiation of SSI, where there is strong evidence of introgression into the root of the radiation and substantial evidence of secondary adaptive introgression into both trophic specialists from at least three different biogeographic regions of the Caribbean and Atlantic coast (26, 31). Lake Chichancanab is an endorheic basin with no above-ground hydrological connections to the ocean and is substantially more geographically isolated than SSI, a small island in the approximate center of the Caribbean. Furthermore, the sister species to the Chichancanab species flock, *C. artifrons*, is abundant along the entire coast of the Yucatán peninsula, suggesting that any gene flow into Chichancanab would likely first have to pass through *C. artifrons* populations. An additional factor is the water chemistry of Lake Chichancanab which is near-saturation with calcite and gypsum and may be a greater physiological barrier to marine pupfish colonists than the slightly hypersaline lakes of SSI (33–36). Thus, geography and ecology play an important role in determining the relative importance of adaptive introgression in radiations.

### De novo mutations primarily drive adaptive divergence of the zooplanktivore

The importance of de novo mutations in adaptive radiations remains contentious, in part due to the time needed for de novo beneficial mutations to arise (11, 15). Interestingly, we found that 30.6% of the zooplanktivore adaptive variants were detected only within Chichancanab species, suggesting their de novo origins within the basin. This proportion is 15.3 times greater than the contribution of de novo mutations to adaptive divergence of the scale-eater in the SSI radiation. A primary factor could be that the estimated effective population size for the Chichancanab radiation is approximately twice that of the SSI radiation (26), indicating that selection on de novo mutations is more effective, with a higher probability of fixation, all else being equal (14). There may also be a lack of existing standing genetic variation within Caribbean pupfish populations that is adaptive for zooplanktivory. In support of this hypothesis, although radiations should proceed along genetic lines of least resistance (69), the primary axis of craniofacial morphological variation along which the zooplanktivore diverged is far removed from the first two major axes of craniofacial variation within Caribbean pupfish species (29). Evolving along the line of greatest resistance instead of least resistance may require de novo mutations. While the SSI scale-eater is also highly divergent, it is closer to existing craniofacial variation within Caribbean pupfish species and in the direction of greatest variance along the first two principal component axes (29). Similarly, mutational target size for the craniofacial features of a zooplanktivore may be much smaller than the features of a scale-eater, resulting in less standing genetic variation available for adaptation to this trophic niche. Alternatively, we may not have detected these de novo mutations in other populations, but this is unlikely to explain all de novo mutations given our extensive sampling of *Cyprinodon* species across the clade and spanning the Caribbean (Figure 1B). Thus, our results highlight that de novo mutations may play a more important role in driving young adaptive radiations than currently appreciated (12, 13), especially when ecotypes are highly divergent from other species within the radiation.

### Minimal parallelism in the genes underlying adaptive divergence of trophic specialists

We found only two candidate genes (*DNM1* and *SPTLC3*) among the 55 zooplanktivore candidate genes in the Chichancanab radiation that overlapped with the hundreds of candidate genes identified so far in the SSI radiation (26, 28, 30, 32, 42–46). Consistent with trophic specialization requiring the pursuit of mobile prey, both genes are involved in motor neuron development. *DNM1* is a dynamin with nervous tissue-specific expression that is crucial for the proper formation of motor neuron axons (47). *SPTLC3* is involved in the biosynthesis of sphingolipids (70), and suppression of *SPTLC3* in zebrafish results in motor neuron axon defects (48).

The extremely limited number of shared adaptive genes is surprising given the high conservation of gene function in both morphological development (71, 72) and behavior (73), and the frequent parallelism of candidate genes across vertebrate adaptive radiations, particularly for craniofacial genes. For example, *BMP4* is involved in diversification in both Galápagos finches (74) and African cichlids (75, 76). Furthermore, within the SSI radiation, many of the same genes show parallel patterns of differential expression even within divergent trophic specialists (26, 43). Nonetheless, parallelism of shared regulatory networks, rather than specific genes, is still a possibility.

### Rapid divergence in tooth morphology of the zooplanktivore through WNT10A

Another reason for non-parallelism is that trophic specialists in each radiation have diverged in different sets of traits during adaptation to different ecological niches (29). Specifically, the Chichancanab radiation has diversified in tooth size over 130 times faster than background rates, driven by the extremely short and narrow teeth of the zooplanktivore (Figure S1; (29). We identified a well-characterized tooth development gene in the Wnt pathway, *WNT10A*, with many de novo mutations in flanking (*n* = 2) and intronic (*n* = 13) regions. Mutations in *WNT10A* are associated with tooth developmental disorders in humans (62) and both knockout and overexpression experiments in sticklebacks and zebrafish have demonstrated *WNT10A*‘s role in teeth development and regeneration (61, 63). Consistent with *WNT10A* playing a role in rapid divergence of tooth morphology, we found that inhibiting the Wnt pathway in the *C. beltrani* generalist during pre-metamorphosis development resulted in narrower teeth, similar to the reduced tooth sizes in the zooplanktivore *C. simus* (Figure 5C). This indicates that the substantial number of de novo mutations in *WNT10A* may be suppressing the Wnt pathway, resulting in a substantial reduction in tooth size in the zooplanktivore.

### Increased visual acuity in the zooplantkivore is consistent with unique selection on vision-related genes

One of the more striking features of both *Cyprinodon* radiations is the evolution of novel trophic specialists from a generalist ancestor: the scale-eater (*C. desquamator*) in the SSI radiation and the zooplanktivore (*C. simus*) in the Chichancanab radiation. Although these are distinct dietary specializations, they share a similarity in that they require tracking free-swimming prey. Indeed, four of the top 15 enriched GO terms of candidate genes in the SSI scale-eater were related to eye development (26). Similarly, several of the candidate genes we identified in the zooplanktivore were related to eye development, including *ASIC3*, *MAB21L2*, and *CRIM1*. *CRIM1* is the most promising candidate because it is one of three candidate genes with multiple nonsynonymous mutations (Figure 2). In zebrafish, a two-base pair deletion in *CRIM1* resulted in abnormal lenses as well as a large eye-to-head ratio (54). Interestingly, one of the defining morphological features of *C. simus* is its large eye-to-head ratio (33) (Figure 1A). Consistent with candidate genes related to eye development, we confirmed that *C. simus* exhibits a much stronger optomotor response than *C. beltrani* (Figure 5B), indicating that this specialist has greatly improved visual acuity, even when reared in a common laboratory environment.

Although both radiations contained many vision-related candidate genes, selection on opsin genes was unique to Chichancanab. Most notably, we found several adaptive variants associated with *SWS2* (blue-sensitive), and *OPN1LW* (red-sensitive) in the zooplanktivore. Interestingly, both opsins contain many candidate de novo mutations, and *SWS2* is the youngest selective sweep in the zooplanktivore. One explanation could be that most Caribbean *Cyprinodon* species are benthic and live in similar habitats, resulting in limited standing genetic variation in opsins. Additionally, most environments (including SSI and Chichancanab) are relatively shallow, so differences in depth—a common selective pressure on opsin evolution (77)—are likely not a strong selective force on SSI. Thus, the main selective force acting on vision in Chichancanab may be prey capture efficiency. In many fish species, zooplanktivory is associated with higher expression of shorter wavelength opsins (e.g., *SWS1* (77)), whereas piscivory is not often associated with changes in opsin expression (78, 79).

### Divergent sperm morphology in Chichancanab species is consistent with selection on spermatogenesis genesis

Although the role that premating sexual selection plays in speciation has historically been the main focus (80, 81), there is both theoretical and empirical support for the role of postmating sexual selection in maintaining reproductive isolation via postmating prezygotic reproductive barriers (82–87). In the SSI radiation, species differ in pigmentation, and a handful of the candidate adaptive variants fall in regulatory regions of genes associated with pigmentation (26), but not spermatogenesis. Furthermore, there is evidence for premating isolation by female preference in the scale-eater (88). However, in Chichancanab, the two species tested in our data set (*C. labiosus* and *C. beltrani*) were not found to visually distinguish between different species in the radiation in laboratory mate choice trials (89).

Unlike the SSI radiation, we did not find any candidate adaptive variants associated with pigmentation in Chichancanab; instead, we identified adaptive variants associated with four spermatogenesis genes, *ERC2*, *TBC1D25*, *C4orf22*, and *STK35*. Consistent with previous work demonstrating the importance of these genes in spermatogenesis (57–60), we found that *C. simus* does indeed have larger sperm heads than *C. beltrani*, suggesting a unique role for post-mating prezygotic reproductive barriers in this radiation (Figure 5D).

### Parallelism in the stages of adaptation: refinement with de novo and nonsynonymous mutations

Adaptation may occur in stages, with less pleiotropic cis-regulatory changes occurring before potentially more pleiotropic amino acid changes (26, 90, 91). Selection is also predicted to act on standing variation first (14, 92). In both radiations, we observed strong evidence for a refinement stage in which coding substitutions were selected on last. In Chichancanab, these late-sweeping coding substitutions were also de novo mutations (Figure 4). The simplest explanation for parallelism in more recent ages of adaptive de novo mutations is the waiting time required for beneficial mutations to (1) arise and (2) reach a substantial frequency within the population (14, 92). These variants may only be selected in the later stages of an adaptive walk because nonsynonymous mutations are often pleiotropic and likely deleterious in the ancestral genetic background (14, 93). Thus, they may not be advantageous until later in the adaptive walk, when sign epistasis may reverse negative pleiotropic effects (94–98). Our results highlight that this final refinement stage is likely a general pattern during adaptive divergence.

## Materials and Methods

### Sampling and genotyping

In 2022, we collected Lake Chichancanab specimens (*C. labiosus*, *C. beltrani*, and *C. simus*) and *C. artifrons* (Figure 1C). Additional species were collected from the wild or sourced from captive breeding populations in previous years. All specimens were euthanized in an overdose of buffered MS-222 (Fiquel, Inc.) following approved animal care and use protocols from the University of California, Berkeley and the University of California, Davis. All specimens used in this study are catalogued in the Museum of Vertebrate Zoology Fishes collection (MVZ:Fish:1410-1499).

Individual DNA samples were sequenced using Illumina HiSeq 4000 and NovaSeq. We mapped reads to the UCB_CyNevMio_1.0 *Cyprindon nevadensis mionectes* reference genome (GCA_030533455.1). Variants were called using *GATK* (v.4.5.0.0) and filtered according to best practices (99), resulting in 23,960,536 SNPs. See SI for more details.

### Population structure and demographic history of Yucatán Cyprinodon pupfishes

To assess population structure, we first pruned SNPs in linkage disequilibrium using *PLINK* (v. 1.9), resulting in 5,476,855 SNPs. With this linkage disequilibrium pruned data set, we visualized population structure with a PCA (*PLINK*) and *ADMIXTURE* (v1.3) (100).

We estimated demographic history using *MSMC2* (101) using three individuals with the highest mean depth. See SI for more details.

### Identifying and characterizing candidate genes

We calculated *F_st_* genome-wide, per site, and in non-overlapping 10-kb windows in *VCFtools* (v0.1.16) for all pairwise combinations of Yucatán *Cyprinodon* species (102). With the 10-kb windowed *F_st_* values, we calculated a modified, normalized population branch statistic (PBS), *PBS_nj_* (103). We classified SNPs as candidate adaptive variants if they were fixed or nearly fixed (per-site *F_st_* >0.95) compared to other species within the radiation and were among the top 1% outliers out of all *PBS_nj_* windows, following the threshold used in (41).

We classified variants within 20 kb of a gene (upstream or downstream) as flanking. For variants within a gene, we further classified them into intronic, synonymous, or nonsynonymous using *SnpEff* (104). To characterize the origin of candidate variants, we extracted the alleles from every species for each candidate variant. We classified variants that were found in any other populations outside of Lake Chichancanab as standing genetic variation. We further subdivided this group into variants that were found in other species but not in *C. artifrons*. Finally, if the alternative variant was only detected within Chichancanab species, we classified it as de novo. See SI for more details.

### Introgression

We tested for evidence of introgression in Yucatán *Cyprinodon* species using *Dsuite* with *Cualac tessellatus* as the outgroup, calculating *f_4_*-ratios, *D* statistics, and *f*-branch statistics (49). For species trios that showed evidence of introgression where *C. simus* was either P1or P2, we then looked for introgression regions across the genome by calculating the distance fraction (*d_f_* (50)), in non-overlapping windows of 91 informative SNPs. To estimate introgression block sizes, we merged neighboring windows that were introgression outliers and summed the total size of the adjacent block. See SI for more details.

### Timing of selective sweeps

For each candidate gene, we used *starTMRCA* (51) to estimate the age of selective sweeps, extracting a 1-Mb region surrounding the candidate variant (51). We used a fixed recombination rate of 2x10^-8^ (swordtail fishes; (105)) and a mutation rate of 1.56 x 10^-8^ substitutions per base pair (Caribbean *Cyprinodon* species; 26). We then conducted separate generalized linear models to test if the sweep age was correlated with the proportion of nonsynonymous SNPs, proportion of de novo SNPs, and the number of SNPs. See SI for more details.

### Optomotor response, sperm morphology, and Wnt inhibition experiment

We tested for differences in visual acuity between the generalist/detritivore, *C. beltrani*, and the zooplanktivore, *C. simus*, by conducting an optomotor response behavioral assay (56) on lab-reared *C. beltrani* (F1; *n* = 7) and lab-reared *C. simus* (F10+; *n* = 5). Because the data were nonparametric, we used a Wilcoxon rank-sum test to assess differences in optomotor response.

To test for differences in sperm morphology, we collected sperm samples from five individuals per species in the lab (F10+*C. simus,* F1 *C. artifrons,* F1 *C. beltrani*). After fixing the sperm sample in a 4% PFA solution stained with Rose Bengal, we plated and imaged sperm under oil immersion with a 63x objective. We measured 10-30 sperm per individual in ImageJ (106). We fit separate linear mixed-effects models for each trait of interest (sperm head, midpiece, and flagellum lengths) with species as a fixed effect and a random intercept for individual ID.

To confirm the role of the Wnt pathway in proper jaw and teeth development, we conducted an experiment chemically inhibiting the Wnt pathway using iCRT14 during metamorphosis (8dpf to 22 dpf) in *C. beltrani*. To control the exact time of development, we first performed in vitro fertilizations and incubated developing embryos at ∼26 °C until hatching (∼8dpf). On the day of hatching, we haphazardly split broods into control (DMSO) and experimental groups (100 nanomolar iCRT14) to control for batch and family effects. After euthanizing and staining fish with alizarin red (bone) and alcian blue (cartilage; (107)), we quantified the number of ossified teeth and measured the tooth length and base width for the left and right teeth closest to the mandibular symphysis using ImageJ (106). We fit separate linear mixed-effects models for average tooth length and base width with experimental treatment as a fixed effect and a random intercept for brood/replicate. For the number of ossified teeth, we used a generalized linear mixed-effect model with a Poisson family and log-link function. See SI for more details.

## Acknowledgments

MCK was supported by a Miller Postdoctoral Research Fellowship from the Miller Institute for Basic Research in Science, University of California Berkeley. This research was funded by NSF CAREER 1749764 and NIH 5R01DE027052-02 awards to CHM. We thank the Gerace Research Center and Troy Day for logistical support in the Bahamas, and the governments of Mexico and the Bahamas BEST commission for permission to collect and export samples (Permit No. PPF/DGOPA-001/20). All research procedures and animal care protocols (AUP-2021-02-14062-1 and AUP-2021-07-14515) were approved by the University of California, Berkeley Animal Care and Use committee and the UC Davis Institutional Animal Care and Use committee. Specimens are deposited in the Museum of Vertebrate Zoology (MVZ:Fish:1410-1499).

## Supporting Information

**The supporting information includes**

Supporting text (SI methods)

Tables S1 to S4

Figures S1 to S10

SI References

## SI Methods

### Sampling

We collected Lake Chichancanab specimens in 2022 over two days using a 5 x 1.3 m seine net with 1.6 mm mesh size. We sampled two sites, an entrance road by La Presumida, and a bridge crossing the narrow section of the lake on the road to San Diego (Fig. 1C). *C. labiosus* and *C. beltrani* were sampled from both sites for this study, whereas *C. simus* was only found at La Presumida. *C. artifrons* were collected by cast net from a coastal estuary at Sisal. Additional specimens were collected from the Bahamas (Lake Cunningham, New Providence Island and Crescent Pond, San Salvador Island), the Dominican Republic (Laguna Bavaro), or Fort Fisher, North Carolina as described previously (Martin 2016) or sourced from the American Killifish Association, the London Zoological Society, the Dallas Children’s Aquarium (*Megupsilon aporus*), and Frans Vermuelen (*Cyprinodon dearborni*). All specimens were euthanized in an overdose of buffered MS-222 (Fiquel, Inc.) following approved animal care and use protocols from the University of California, Berkeley and the University of California, Davis. All specimens used in this study are catalogued in the Museum of Vertebrate Zoology Fishes collection (MVZ:Fish:1410-1499).

The London Zoological Society provided our live colony of *C. simus* in 2015, originating from a much earlier field collection, which we subsequently raised for ten generations. We collected live colonies of *C. beltrani* and *C. artifrons* in 2024 from La Presumida, Lake Chichancanab and Sisal, respectively, over four days using a 5 x 1.3 m seine net with 1.6 mm mesh size.

### Sequencing, Variant calling and filtering

Individual DNA samples were resequenced using Illumina Hiseq 4000 and Novaseq. Raw reads were trimmed with *fastp* (v0.23.4) (1). Reads were mapped to the UCB_CyNevMio_1.0 *Cyprindon nevadensis mionectes* reference genome (GCA_030533455.1) with *bwa mem* (v0.7.17) (2). We followed the *GATK* (v.4.5.0.0) genotyping pipeline to call variants (119). Duplicate reads in the bam files were marked with *Picard MarkDuplicates* (GATK v4.5.0.0) (118). Coverage and mapping quality were assessed with *qualimap* (v2.2.2) (4). We used *HaplotypeCaller* (-ERC GVCF) v4.5.0.0 to call variants for each individual and stored variants in the GenomicsDB datastore format prior to using *GenotypeGVCFs* to perform joint genotyping. We restricted our analyses to biallelic SNPs and applied the recommended *GATK* hard filters (QD < 2, QUAL < 30, SOR > 3, FS > 60, MQ < 40, MQRankSum < -12.5, ReadPosRankSum < -8) to filter SNPs (118, 120). Missing genotypes were reset to ./. from 0/0 due to how *GATK* represents missing genotypes in version 4.5.0.0 (7).

After hard-filtering and variant calling, we filtered the data set further by removing SNPs with minor allele frequency < 0.05, more than 10% missing, depth less than 10x, and GQ less than 20 using *BCFtools* (v1.16)(8). This resulted in a final data set that included 23,960,536 SNPs.

### Population structure of Yucatán Cyprinodon pupfishes

To assess population structure, we first pruned SNPs in linkage disequilibrium using *PLINK* (v. 1.9) with the following parameters: “-indep-pairwise 50 5 0.2” (9). This filtered our data set from 23,960,536 SNPs to 5,476,855 SNPs. With this linkage disequilibrium pruned data set, we first filtered the full VCF file to only include relevant Yucatán *Cyprinodon* species (i.e., *C. artifrons, C. beltrani, C. labiosus,* and *C. simus*), then conducted a PCA using *PLINK*. Next, we used *ADMIXTURE* (v1.3) to determine the optimal number of population clusters that best fit the data and to assign individuals to their corresponding population clusters (10). We performed this analysis and calculated cross-validation error for the number of clusters K = 1-6. Models in ADMIXTURE with K=2-4 were equally supported (Figure S2A). However, given morphological distinctness and the results of the PCA, we present the results for when K = 4 (Figure 1D).

### Demographic history of Yucatán Cyprinodon pupfish

We estimated demographic history using *MSMC2* (11). For each Yucatán pupfish species, we selected three individuals with the highest mean depth and generated single-sample VCF files, individual mask files, and “mapability” mask files. We then ran *MSMC2* with default parameters except we set phasing to “unphased” and “P_PAR=8*1+25*1+1*2+1*3.” For plotting, we used a generation time of 1 year and a mutation rate of 1.56 x 10^-8^ substitutions per base pair, estimated from lab-reared pedigrees of Caribbean *Cyprinodon* species (12).

### Identifying candidate genes

We calculated *F_st_* genome-wide, per site, and in non-overlapping 10-kb windows using the “--weir-fist-pop” function in *VCFtools*(v0.1.16) for all pairwise combinations of Yucatán *Cyprinodon* species (13). We chose 10-kb windows because linkage disequilibrium substantially decayed at this distance (Figure S3), and these windows allowed us to quantify fine-scale genomic variation. With the 10-kb windowed *F_st_* values, we calculated a modified population branch statistic (PBS) for four taxa, *PBS_nj_* (14). This statistic is useful because it makes no assumptions about species topology or polarization (14). We then normalized *PBS_nj_* by the total tree length, as normalized *PBS* is most effective and specific at identifying selective sweeps (15). We classified SNPs as candidate adaptive variants if they were fixed or nearly fixed (per-site *F_st_* >0.95) compared to other species within the radiation and were among the top 1% outliers out of all *PBS_nj_* windows, following the threshold used in (15).

### Introgression

Using the full VCF file, which contained a wide range of species in the family Cyprinodontidae, we tested for evidence of introgression in Yucatán *Cyprinodon* species using *Dsuite* (16). We first excluded any samples/species that had an average depth < 4. We then calculated all possible trios with a species tree based on the most recent phylogeny (17) with *Cualac tessellatus* as the outgroup. We calculated Z-scores to assess the significance of *D* statistics using 1,000 jackknife blocks and considered a Z-score greater than three as evidence for significant introgression (18). We only focused our primary analysis on species trios where P1-P2 included species within the Lake Chichancanab radiation. We used the other trio calculations to calculate *f*-branch (*f_b_*) statistics to account for correlated *f_4_*-ratio scores and to inform the timing of introgression events (16).

For species trios that showed evidence of introgression where *C. simus* was either P1or P2, we then calculated the distance fraction (*d_f_*), a robust metric of introgression for windowed analyses (19), in non-overlapping windows of 91 informative SNPs. We chose this number of SNPs because it resulted in windows of approximately 10 kb. We considered windows to be introgressed if they were in the top X% of absolute *d_f_* value, where X% was the percentage of estimated genome wide introgression (e.g, *f_4_*-ratio) for that focal group, following (20, 21). To estimate introgression block sizes, we merged neighboring windows that were introgression outliers and summed the total size of the adjacent blocks.

### Characterizing candidate genes

After identifying candidate variants, we then used *Bedtools* (v2.31) to see if variants occurred near a gene ( within 20 kb) or within a gene (22) with a GFF annotation file for the *C. nev. mionectes* reference genome (Tian et al. in prep). We classified variants within 20 kb of a gene (upstream or downstream) as flanking. For SNPs within a gene, we further classified them into intronic, synonymous, or nonsynonymous using *SnpEff* (23).

To characterize the origin of candidate variants, we extracted the alleles from every species for each candidate variant. We classified variants that were found in any other populations outside of Lake Chichancanab as standing genetic variation. We further subdivided this group into variants that were found in other species but not in *C. artifrons*. If variants fell within an introgression window, we classified them as “introgressed.” Finally, if the alternative variant was only detected within Chichancanab species, we classified it as de novo.

### Timing of selective sweeps

To gain insight into the timing of selective sweeps in *C. simus*, we used *starTMRCA* (24). Because many genes contained multiple candidate variants, for this analysis, we selected the variant that was in the median position of each gene. We then extracted a 1-Mb region surrounding that variant and removed all sites that contained missing data. For simplification, we only used the generalist/detritivore, *C. beltrani*, as the “reference” population and the zooplanktivore, *C. simus*, as the “selected” population. We used a fixed recombination rate of 2x10^-8^ (swordtail fishes; (25)) and a mutation rate of 1.56 x 10^-8^ substitutions per base pair (Caribbean *Cyprinodon* species; 26). We then ran five separate Markov chains for 30,000 iterations with a proposal standard deviation of 150 (preliminary analyses changing this parameter had minimal influence on results). We then discarded the first 10,000 iterations of each chain as burn-in.

To test for stages of adaptation, we conducted separate generalized linear models to test if the sweep age was correlated with the proportion of nonsynonymous SNPs, proportion of de novo SNPs, and the number of SNPs. For the generalized linear regressions of proportion of nonsynonymous SNPs and proportion of de novo SNPs, we used a quasi-binomial family to account for overdispersion with a logit link function. For the number of SNPs, we used a quasi-Poisson family with a log link function. To test for the significance of sweep age, we used a likelihood ratio test comparing a model with the estimated sweep age as an effect compared to a null model.

### Optomotor response

We tested for differences in visual acuity between the generalist/detritivore, *C. beltrani*, and the zooplanktivore, *C. simus*, by conducting an optomotor response behavioral assay (26). We tested lab-reared *C. beltrani* (F1, *n* = 7) and lab-reared *C. simus* (long-term laboratory colony exceeding ten generations in the lab, *n* = 5). Each fish was tested consecutively with each control or treatment for 1-minute observation periods, with the control observation occurring first, followed by presentation of all-black (positive control) and black-and-white spinning bars (treatment: acuity trial). The order of presentation for positive control and treatment was alternated in each trial. Fish were placed in a suspended cylindrical plastic container 0.3 m in diameter that remained stationary during the course of the trials. Spinning black-and-white or all-black paper was rotated outside the clear plastic container at ∼90 RPM during each observation period.

Due to the data being nonparametric (count, zero-inflated, unequal variance between groups), we conducted a Wilcoxon rank sum test for the on-banded (all black) and on-banded portions separately. Because we had an a priori hypothesis that *C. simus* would display a stronger optomotor response (greater visual acuity), we calculated *p*-values with a one-tailed test.

### Sperm morphology

To test for differences in sperm morphology, we collected sperm samples from five individuals per species in the lab (*C. simus, C. artifrons, C. beltrani*). We first anesthetized the fish with a solution of MS-222. After the fish was anesthetized, we collected 1μL of milt (fish semen) with a 1μL glass microcapillary tube and fixed the sample overnight in a 4% PFA solution stained with Rose Bengal. We then plated sperm on a positively charged slide with a sealed coverslip. We imaged sperm with oil immersion with a 63x objective and measured 10-30 sperm per individual in ImageJ (27). Due to poor sperm samples from some individuals (not sufficient quantities), we ended up with the following sample sizes: three *simus*, four *artifrons*, and five *beltrani*.

To retain information about variation within individuals, we fit separate linear mixed-effects models for each trait of interest (sperm head, midpiece, and flagellum lengths) with population as a fixed effect and a random intercept for individual ID, using *LME4* (28). To assess the significance of the model, we used a Type II ANOVA with a Wald *F* test using Kenward-Roger degrees of freedom. If there was a significant effect of population, we conducted post hoc pairwise tests using the *Emmeans* package (29) and corrected for multiple comparisons using the Tukey method. We did not correct for phylogeny due to how closely related the species are, the limited sample size, and the fact that the only significant differences were between sister species.

### WNT experiment

To confirm the role of the WNT pathway in proper jaw and teeth development, we conducted an experiment chemically inhibiting the WNT pathway using iCRT14 during metamorphosis in *C. beltrani*. To control the exact time of development, we first performed in vitro fertilizations and incubated developing embryos at ∼26 until hatching (∼8dpf). On the day of hatching, we haphazardly split broods into control and experimental groups to control for batch and family effects. We raised experimental groups in tank water with a 100 nanomolar concentration of iCRT14 and control groups in a 100 nanomolar concentration of DMSO (solution used to dilute stock iCRT14). We ran the experiment for two weeks until individuals were 22dpf, changing the solution every 3-4 days. We performed this experiment on four separate broods (∼12 individuals/treatment). One brood was excluded due to improper development (no ossification of the spine) in the controls.

After the experiment, we euthanized fish with MS-222 and stained fish with alizarin red (bone) and alcian blue (cartilage) following a modified published protocol (30). After staining, we prepared whole mounts and imaged fish with fluorescent microscopy (Zeiss LSM880 FCS) to quantify ossification of the jaw and teeth (31). We then dissected out the lower jaw and prepared flat mounts to measure teeth morphology. Specifically, we measured the tooth length and base width for the left and right teeth closest to the mandibular symphysis using ImageJ (27). For statistical analyses, we took the mean of the left and right tooth length and base width.

We fit separate linear mixed-effects models for tooth length and base width with experimental treatment as a fixed effect and a random intercept for brood/replicate, using *LME4* (28). For the number of ossified teeth, we used a generalized linear mixed-effect model with a Poisson family and log-link function. To assess the significance of these models, we used a likelihood ratio test comparing it to a null model without treatment as a fixed effect.

## SI Tables

**Table S1.** Mean genome-wide pairwise *F_ST_* statistics within the Chichancanab radiation and with sister outgroup (*artifrons*).

|  | *labiosus* | *beltrani* | *artifrons* |
| --- | --- | --- | --- |
| *simus* | 0.090 | 0.068 | 0.182 |
| *labiosus* |  | 0.032 | 0.184 |
| *beltrani* |  |  | 0.168 |

**Table S2.** *F_ST_* statistics within the Chichancanab radiation.

| Comparison | $F_{ST} = 1$<br>(fixed) | $1 > F_{ST} \geq 0.95$<br>(nearly fixed) | $0.95 > F_{ST} \geq 0.9$ | $0.9 > F_{ST} \geq 0.8$ | Other | Mean<br>$F_{ST}$ |
| --- | --- | --- | --- | --- | --- | --- |
| *simus* vs *beltrani* + *labiosus* | 19 | 1127 | 3304 | 20173 | 10359605 | 0.07 |
| *labiosus* vs *beltrani* + *simus* | 0 | 0 | 3 | 94 | 10384131 | 0.04 |
| *beltrani* vs *simus* + *labiosus* | 0 | 0 | 0 | 13 | 10384215 | 0.02 |

**Table S3.** Fixed SNPs between *C. simus* and *C. beltrani* + *C. labiosus*. Bold rows indicate SNPs within top 1% of PBS windows. Bolded genes only are genes that have regions that fall within the top 1% PBS percentile.

| Position | Gene | Distance to gene | PBS percentile | Annotation |
| --- | --- | --- | --- | --- |
| Chr 1: 5290600 | **STK35** | 12594 | 1.85 |  |
| Chr 1: 5290602 | **STK35** | 12592 | 1.85 | Fertility, eye development and oxidative stress (32). |
| Chr 1: 5290604 | **STK35** | 12590 | 1.85 |  |
| Chr 2: 17498913 | SLC17A8 | Intron | 1.24 | Glutamate neurotransmitter, inner hair cell dysfunction |
| Chr 3: **11687002** | **AGO3** | **Intron** | **0.34** | **RNA regulator** |
| Chr 3: 32286614 | RBMS3 | 14363 | 22.93 |  |
| Chr 3: 32286627 | RBMS3 | 14350 | 22.93 |  |
| Chr 3: 32286638 | RBMS3 | 14339 | 22.93 | Involved in chondrogenic craniofacial development of zebrafish (33). |
| Chr 3: 32286614 | RBMS3 | 14363 | 22.93 |  |
| Chr 11: 27324192 | GRIA3 | Intron | 2.85 |  |
| Chr 11: 27336469 | GRIA3 | 2772 | 3.18 |  |
| Chr 11: 27336472 | GRIA3 | 2775 | 3.18 | Glutamate receptor |
| Chr 11: 27336476 | GRIA3 | 2779 | 3.18 |  |
| Chr 11: **28255536** | **NA** | **3026** | **0.98** | **NA** |
| Chr 11: **28255616** | **NA** | **3106** | **0.98** | **NA** |
| Chr 11: **28256343** | **NA** | **3833** | **0.98** | **NA** |
| Chr 11: **28257367** | **NA** | **4857** | **0.98** | **NA** |
| Chr 11: 28260047 | NA | 7537 | 2.87 | NA |
| Chr 12: 13854139 | **C4orf22** | 11903 | 1.30 | Spermatogenesis, cilia, flagellum (34) |
| Chr 24: **12693158** | **KDM6A** | **Intron** | **0.05** | **Kabuki syndrome (craniofacial development; 35)** |

**Table S4.** Summary statistics of the distance between closest adaptive candidate SNPs within the same gene.

| Gene | Median Distance | Mean Distance | SD |  |
| --- | --- | --- | --- | --- |
| TM6SF2 | 7430 | 7430 | NA | 2 |
| C19orf10 | 4789 | 4789 | NA | 2 |
| KDM6A | 2892.5 | 2892.5 | 2410.52702 | 3 |
| KCNB2 | 1803 | 1738.75 | 1614.92299 | 5 |
| UBE2T | 1742 | 1742 | NA | 2 |
| TMEM230 | 1734 | 3039 | 3599.52427 | 4 |
| ABCA1 | 1037.5 | 1097.5 | 959.375665 | 5 |
| HAPLN4 | 974 | 1608.21739 | 9927.05273 | 24 |
| GRIN2A | 904 | 1270.6667 | 4273.38008 | 4 |
| DPH1 | 608 | 1224.5 | 1999.25934 | 7 |
| WNT10A | 402 | 1442.17857 | 3986.39626 | 29 |
| OPN1LW | 386.5 | 359.5 | 225.229513 | 5 |
| NIPSNAP3A | 253 | 1451.71429 | 2819.19065 | 8 |
| GALNT15 | 246.5 | 675.166667 | 3156.86746 | 13 |
| SWS2 | 246 | 1381.25 | 2423.03892 | 5 |
| STK35 | 217 | 793.366667 | 1992.8543 | 31 |
| CRIM1 | 201 | 47.829268 | 2176.05381 | 42 |
| PDYN | 187 | 326.371429 | 355.925561 | 36 |
| ADCYAP1R1 | 186.5 | 833 | 1304.35501 | 11 |
| DNM1 | 186.5 | 459.708333 | 677.261781 | 25 |
| DNM1 | 186.5 | 459.708333 | 677.261781 | 25 |
| KCNH8 | 131.5 | 584.366667 | 1030.65532 | 31 |
| TOP1 | 124 | 332.375 | 6687.63393 | 17 |
| AGO3 | 115 | 357.625 | 552.097284 | 9 |
| GMEB2 | 112 | 319.490196 | 2485.05809 | 52 |
| GPNMB | 57 | 57 | 77.781746 | 3 |
| IGFBP4 | 26 | 2254 | 3880.67997 | 4 |
| FAM83D | 23 | 2523.66667 | 4347.74658 | 4 |
| EXOSC3 | 17 | 17 | NA | 2 |
| C4orf22 | 15 | 15 | 9.899495 | 3 |
| TMEM106B | 7 | 11.4 | 10.358571 | 6 |
| THOC3 | 6 | 5.833333 | 3.600926 | 7 |
| MID1 | 1 | 1 | NA | 2 |

## SI Figures

**Figure S1.**
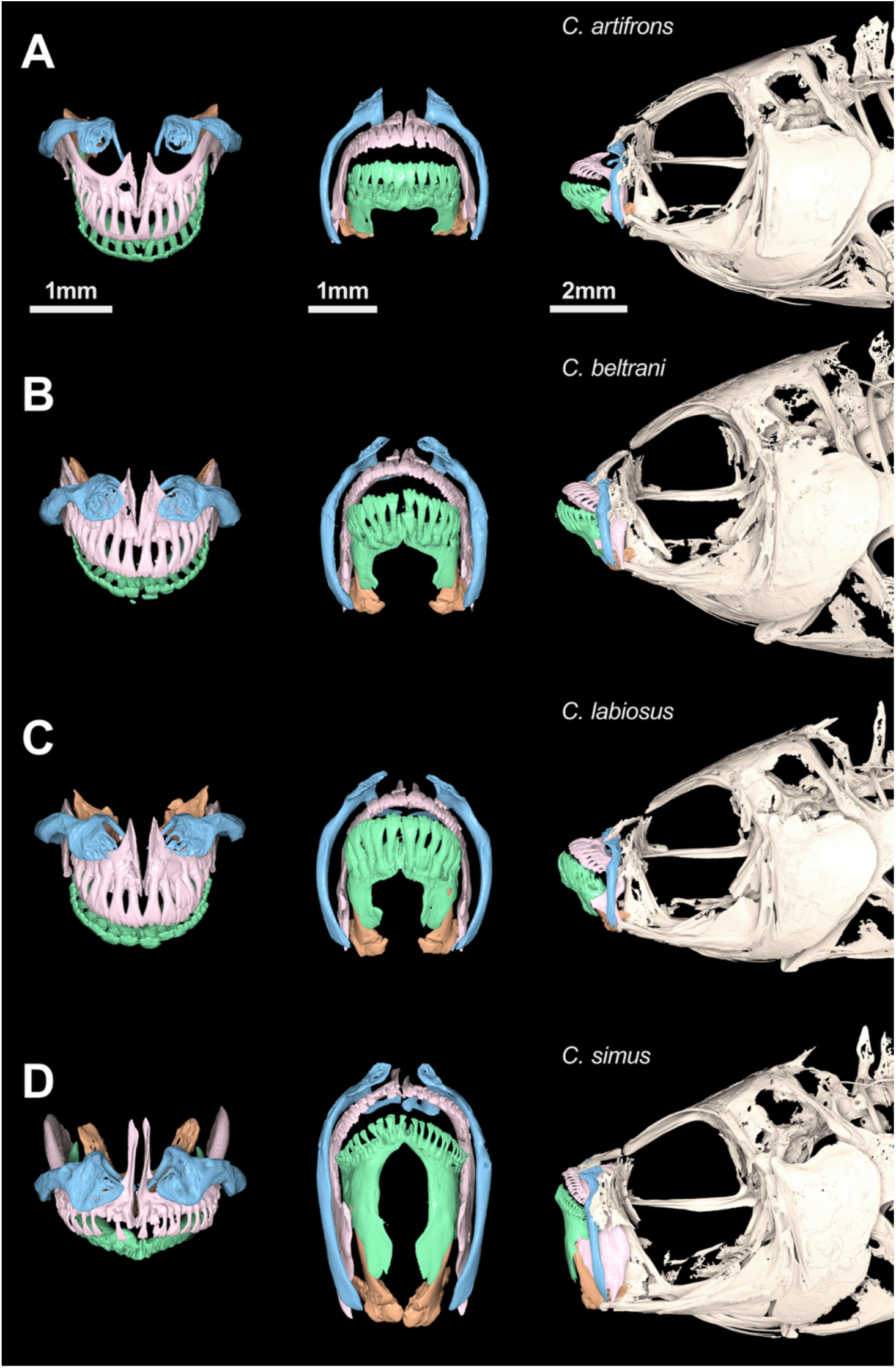
µCT scans of Yucatán pupfish with maxilla colored in blue, premaxilla in pink, dentary in green, and articular in orange.

**Figure S2.**
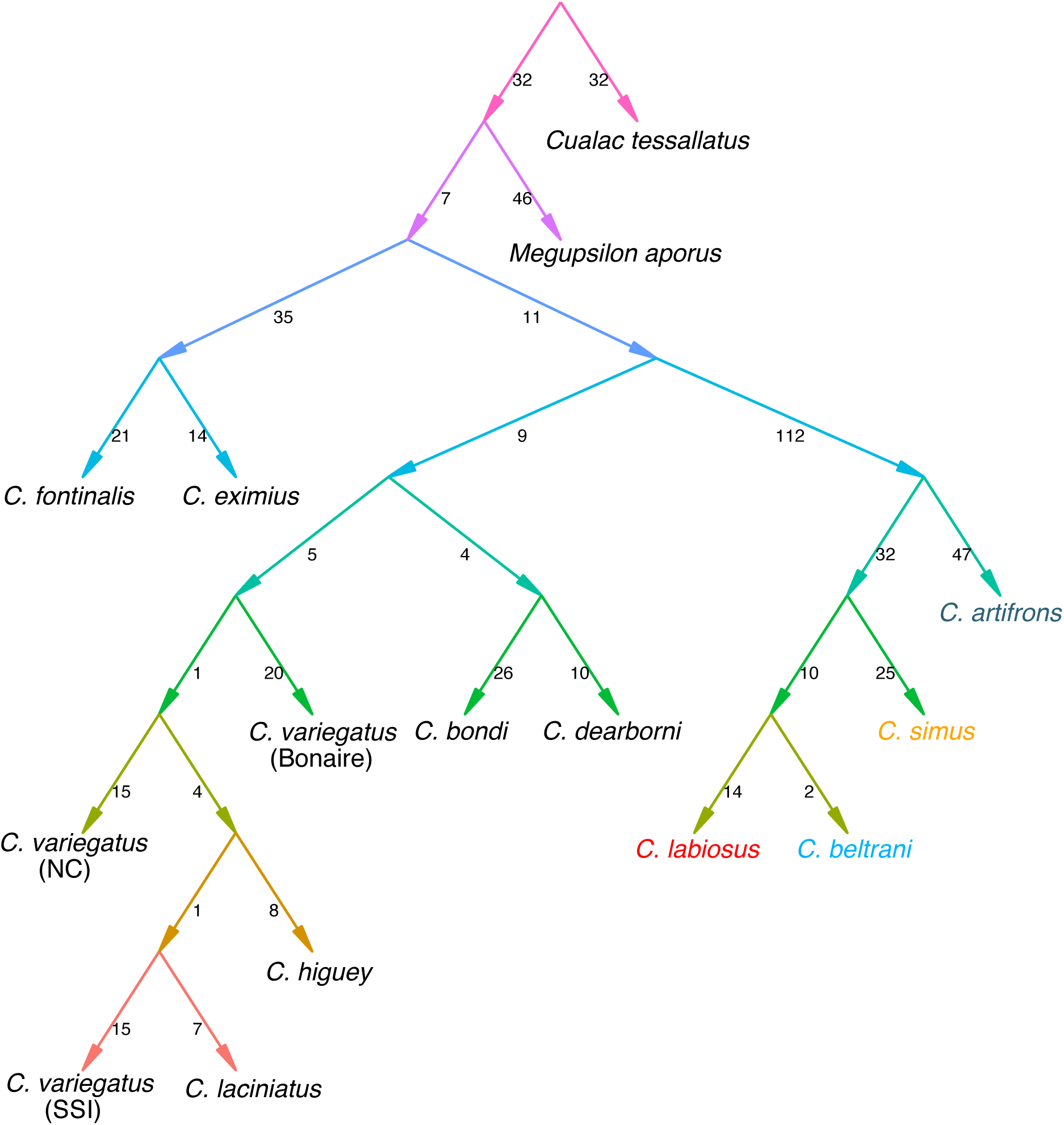
Best fitting admixture graph assuming zero admixture events. Admixture graph was found using the *findGraphs* function in *ADMIXTOOLS2* (36). Number on edges represent drift weights.

**Figure S3.**
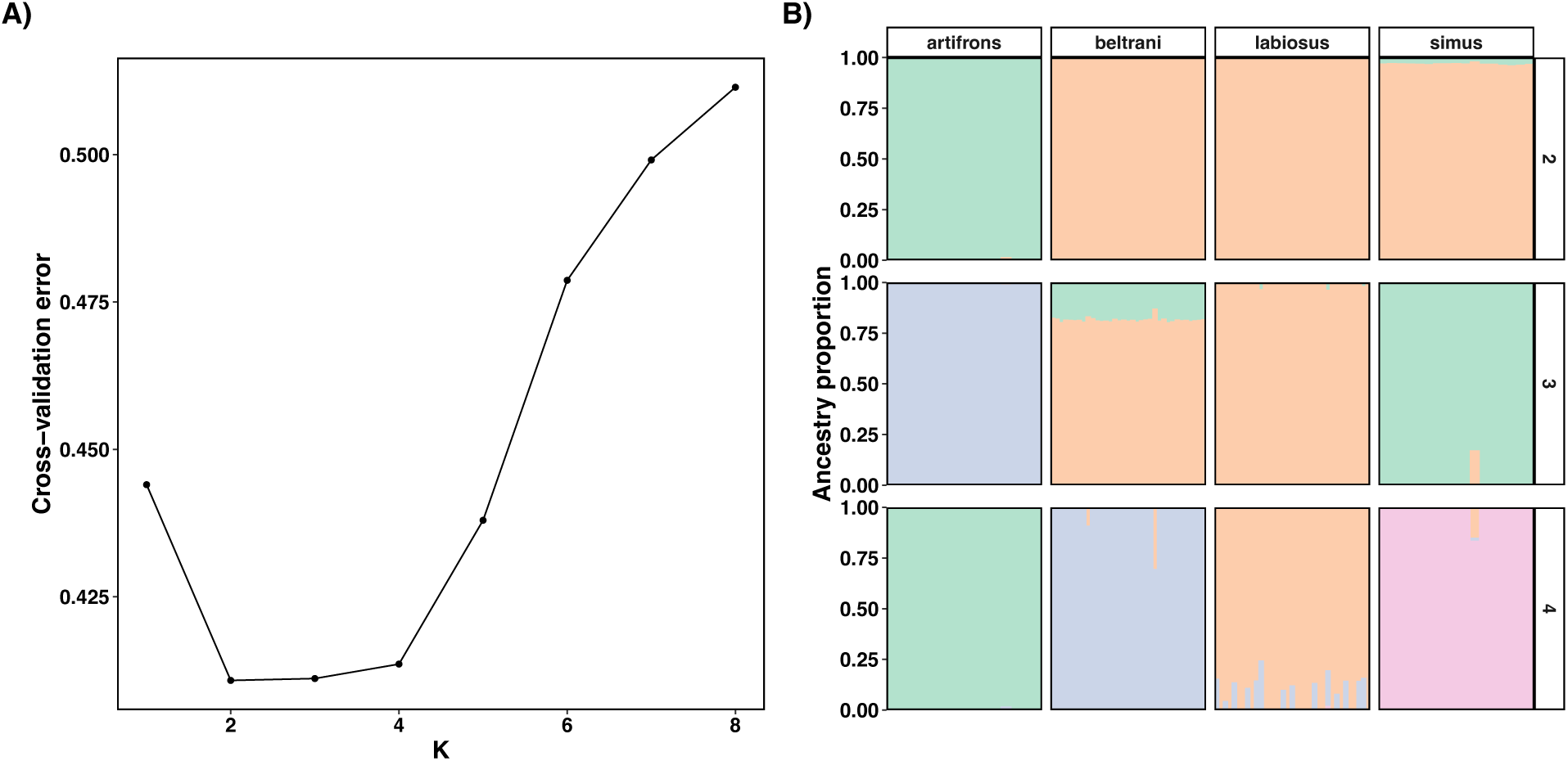
ADMIXTURE results support genetic differentiation of Yucatán Cyprinodon pupfish species. (A) Cross-validation error for different values of K clusters, equally support 2, 3, and 4 clusters. (B) *ADMIXTURE* plots showing the clusters at different values of K.

**Figure S4.**
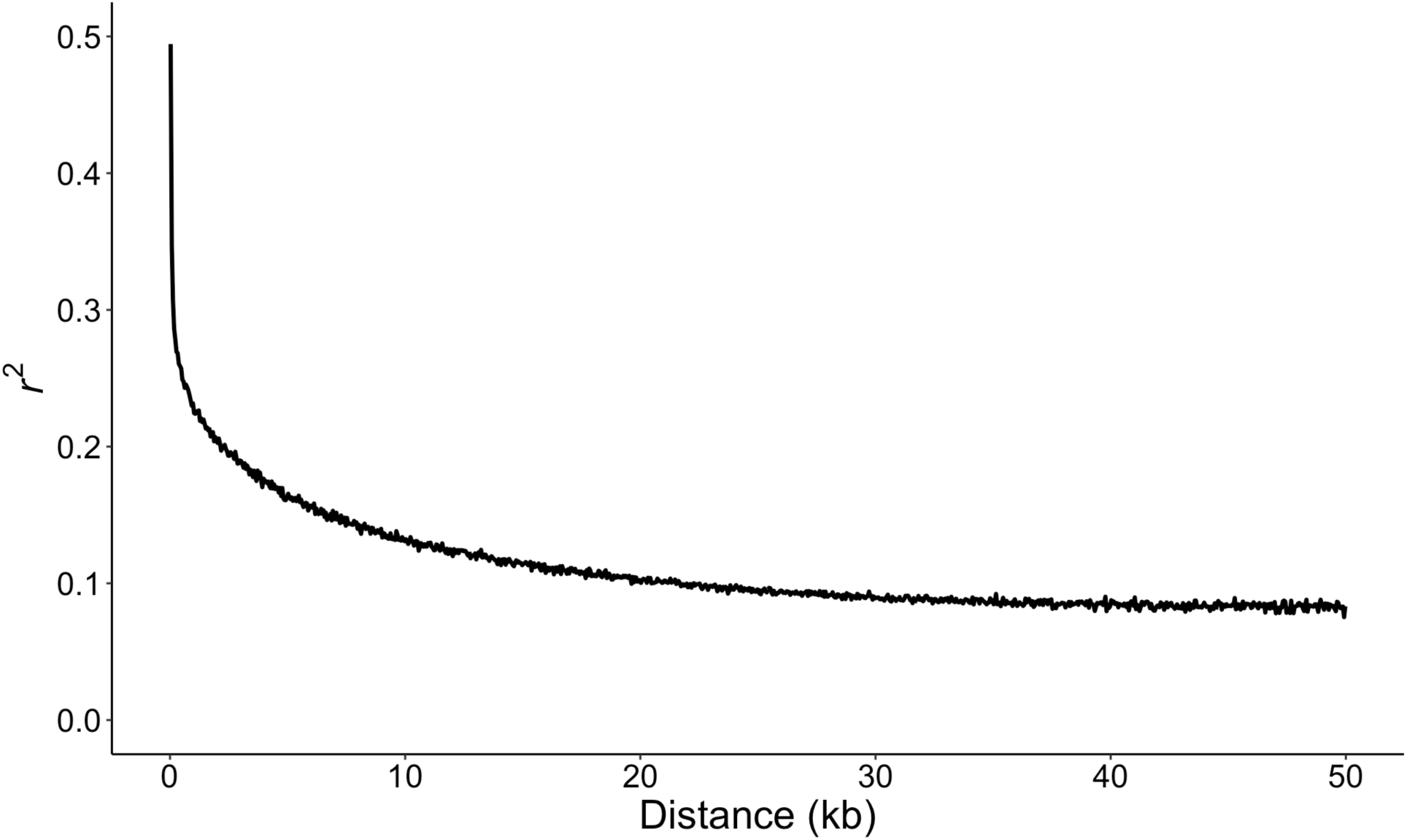
Linkage decay occurs rapidly in *Cyprinodon simus*. Linkage decay was calculated in windows of 1000 kb using *PLINK* (v1.9). The results graphed are a random sample of all calculated *r^2^*values taken for plotting purposes (*n* = 9,645 *r^2^* values).

**Figure S5.**
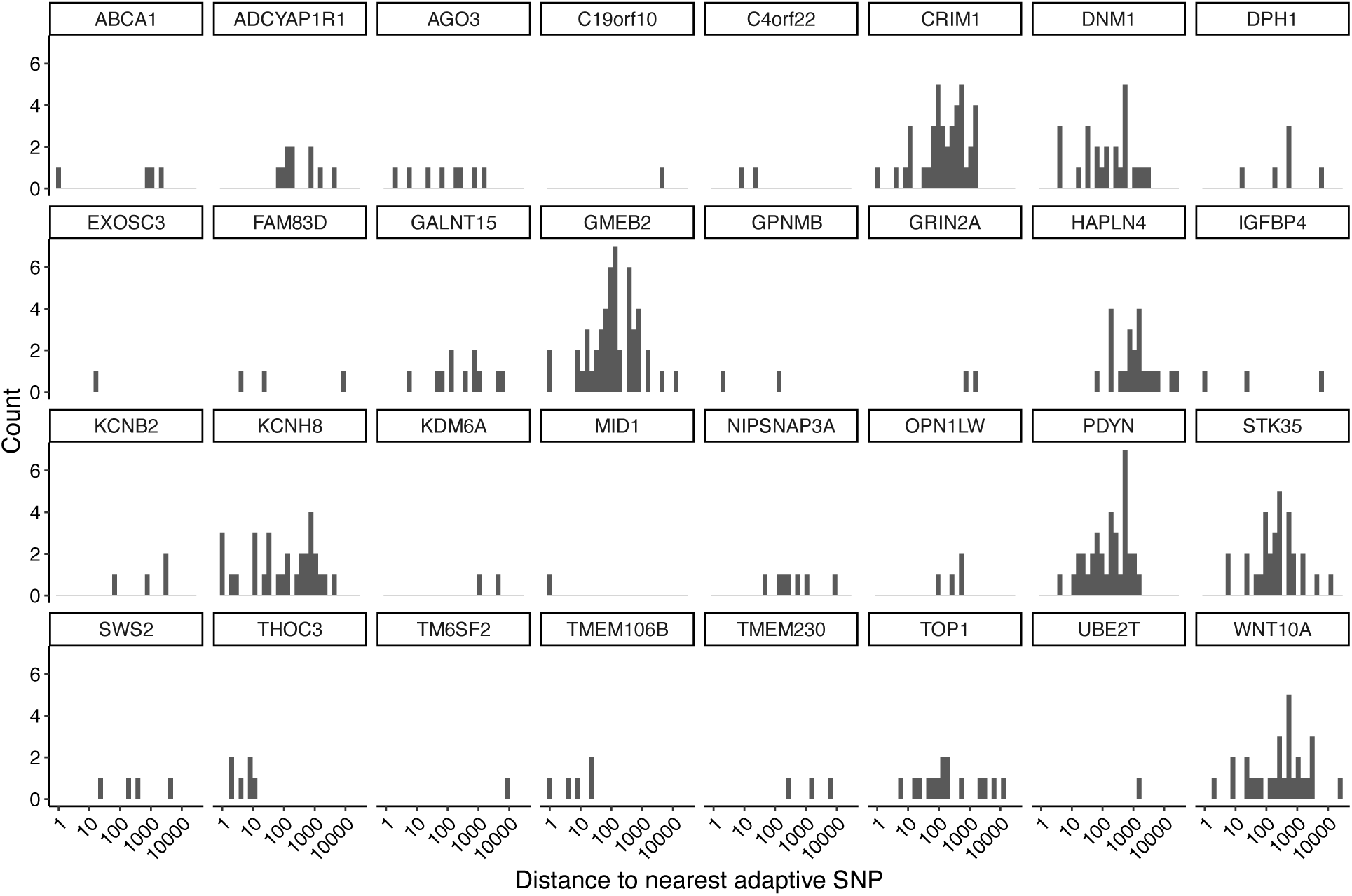
Adaptative SNPs associated with the same gene are generally spaced apart. Distribution of the distance to nearest adaptive SNP within a gene. Note the log scale of the x-axis.

**Figure S6.**
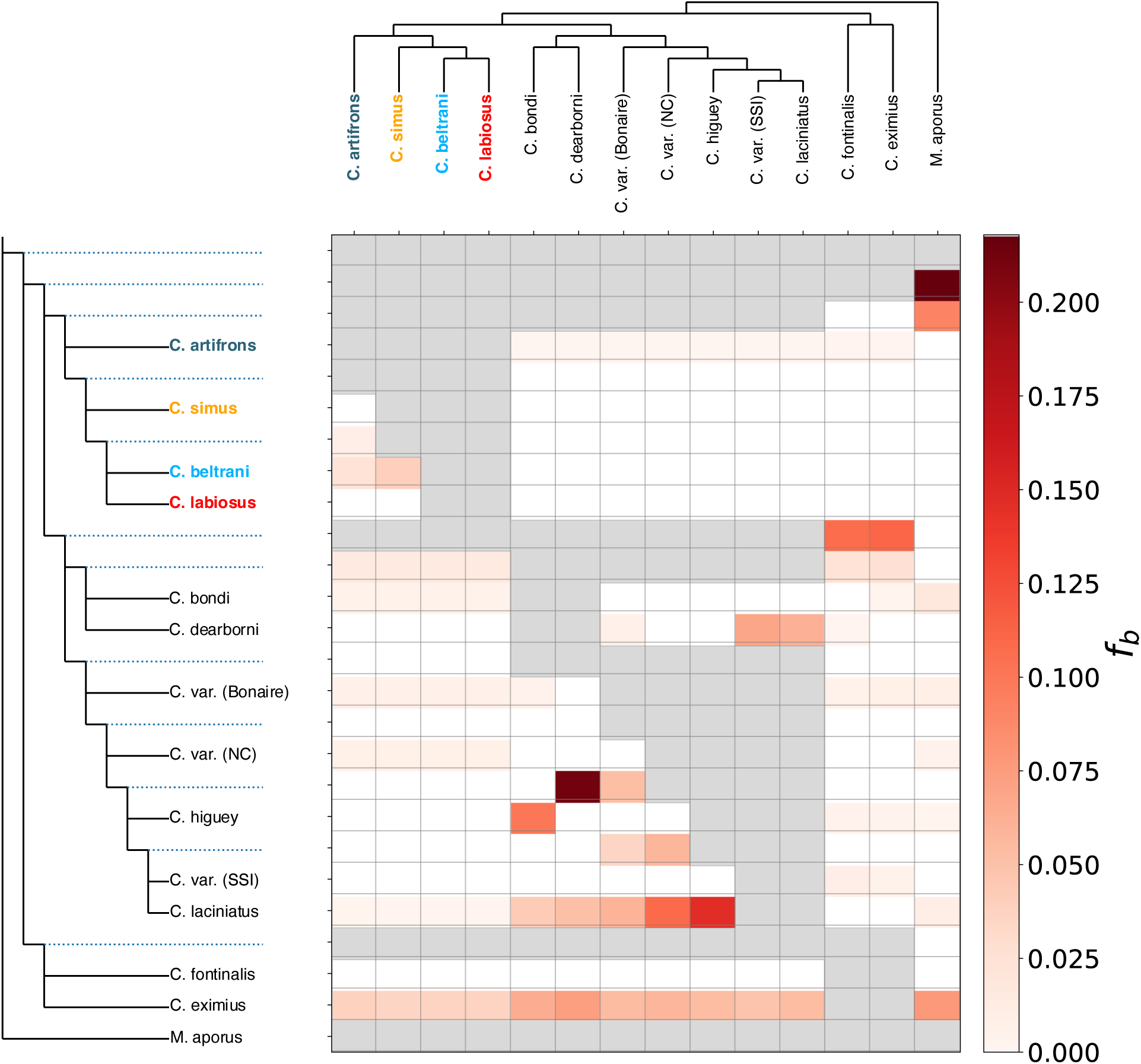
*f-*branch (*f_b_*) show that introgression with desert pupfish occurred before the Chichancanab radiation. Boxes with red color represent significant introgression between P3 species (top) and the species/internal branch on the left. For visualization, non-significant *f_b_* were changed to zero and are completely white boxes. Grey boxes represent comparisons that cannot be made, as *f_b_* cannot be calculated for introgression between sister taxon. Dashed lines on the left are internal branches. SSI: San Salvador Island, Bahamas; NC: North Carolina, USA.

**Figure S7.**
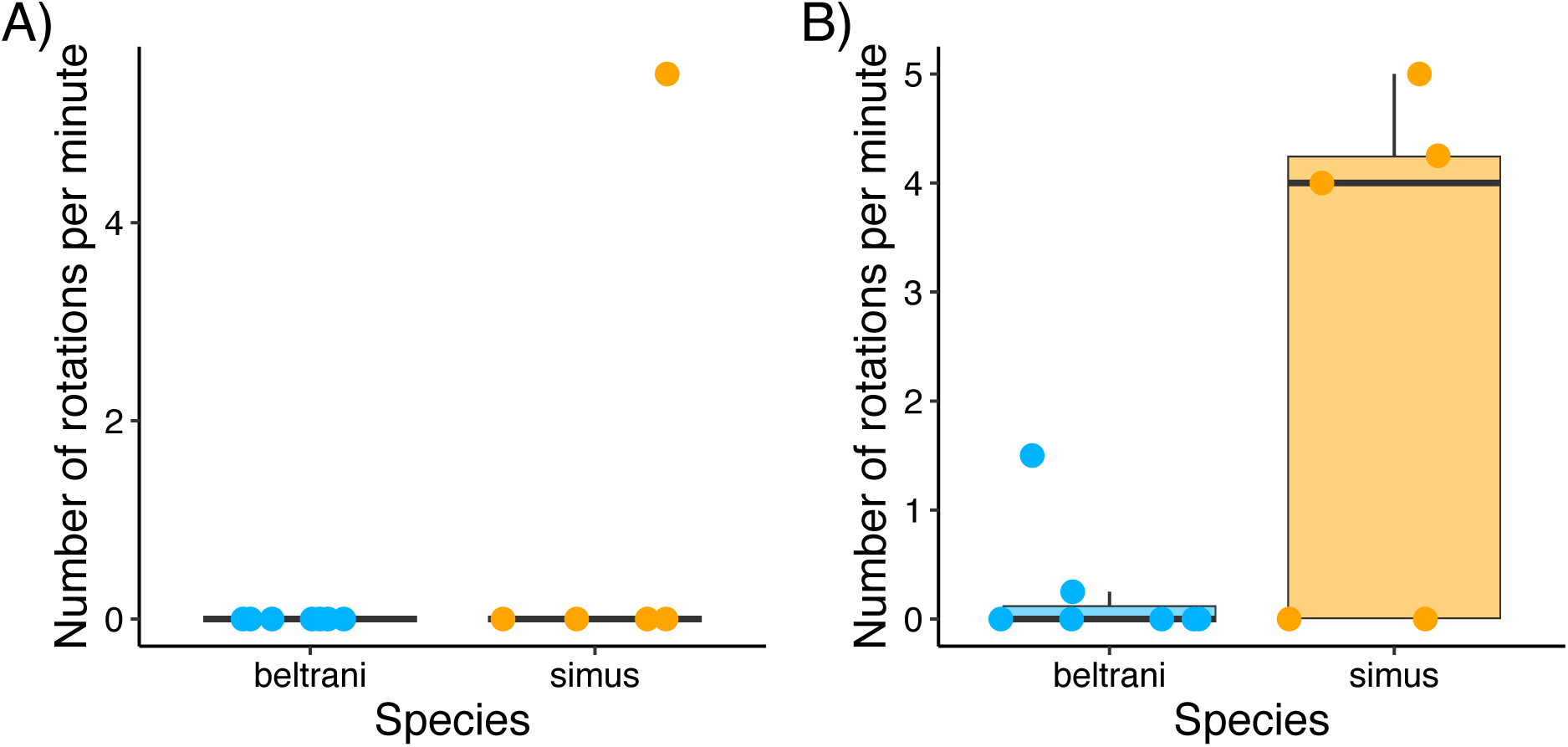
Both species did not have a strong response to either control scenario in the optomotor response experiment. **(A)** Number of complete rotations fish swam when the machine was off and **(B)** when the machine was on but the apparatus was all black. Boxplots and points of the number of complete rotations individuals performed during the “off-banded” portion of the experiment. Note that some boxplots are compressed to a single line because most individuals had no response.

**Figure S8.**
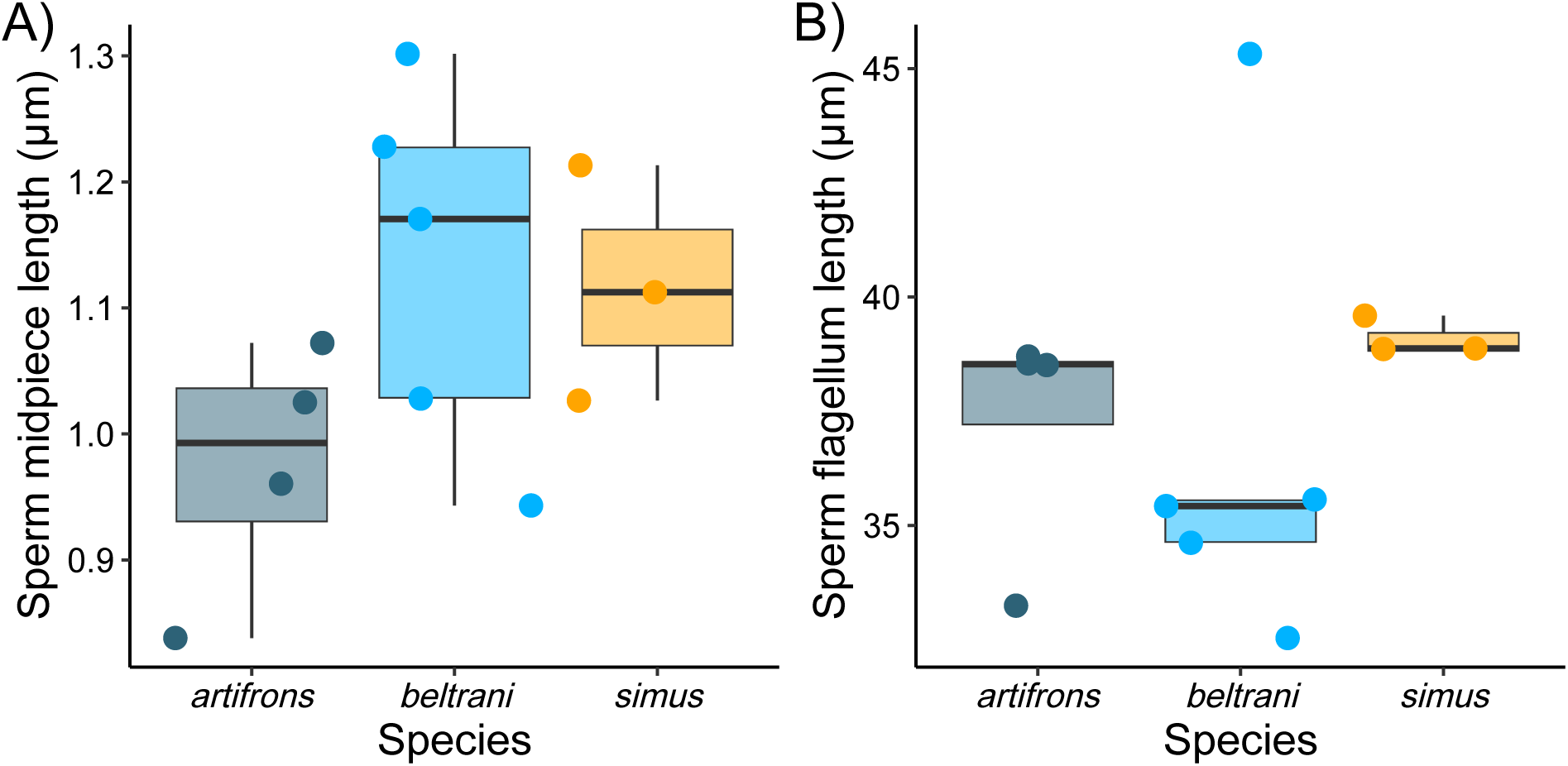
Species do not differ in sperm midpiece size or sperm flagellum length. Boxplots and points of the mean sperm morphological estimate for an individual.

**Figure S9.**
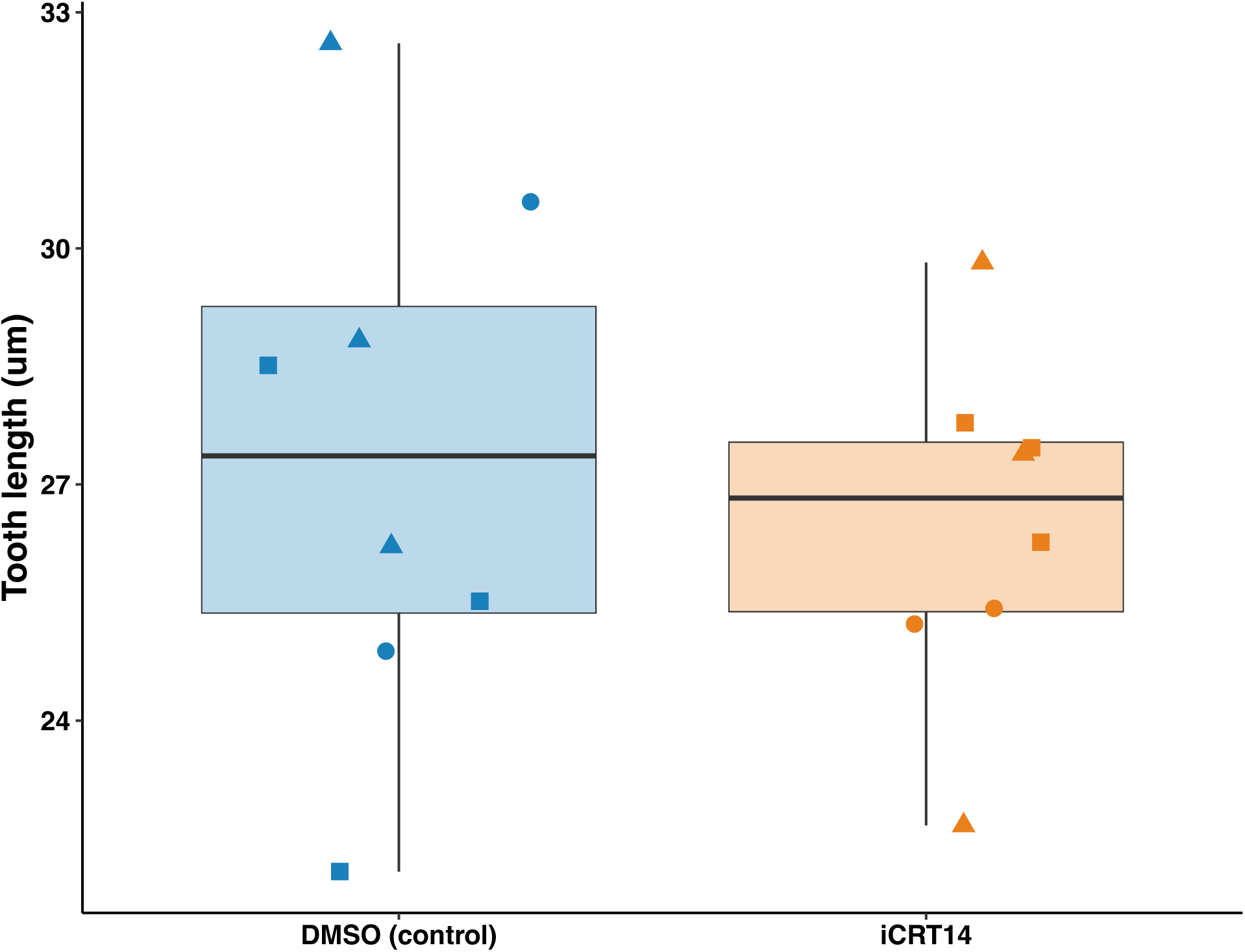
Tooth length was not affected by Wnt inhibition. Boxplot and jittered points of average tooth length of the left and right teeth closest to the mandibular symphysis for *C. beltrani* larvae treated with either a control or Wnt inhibitor (iCRT14). Shapes represent different split-brood replicates.

**Figure S10.**
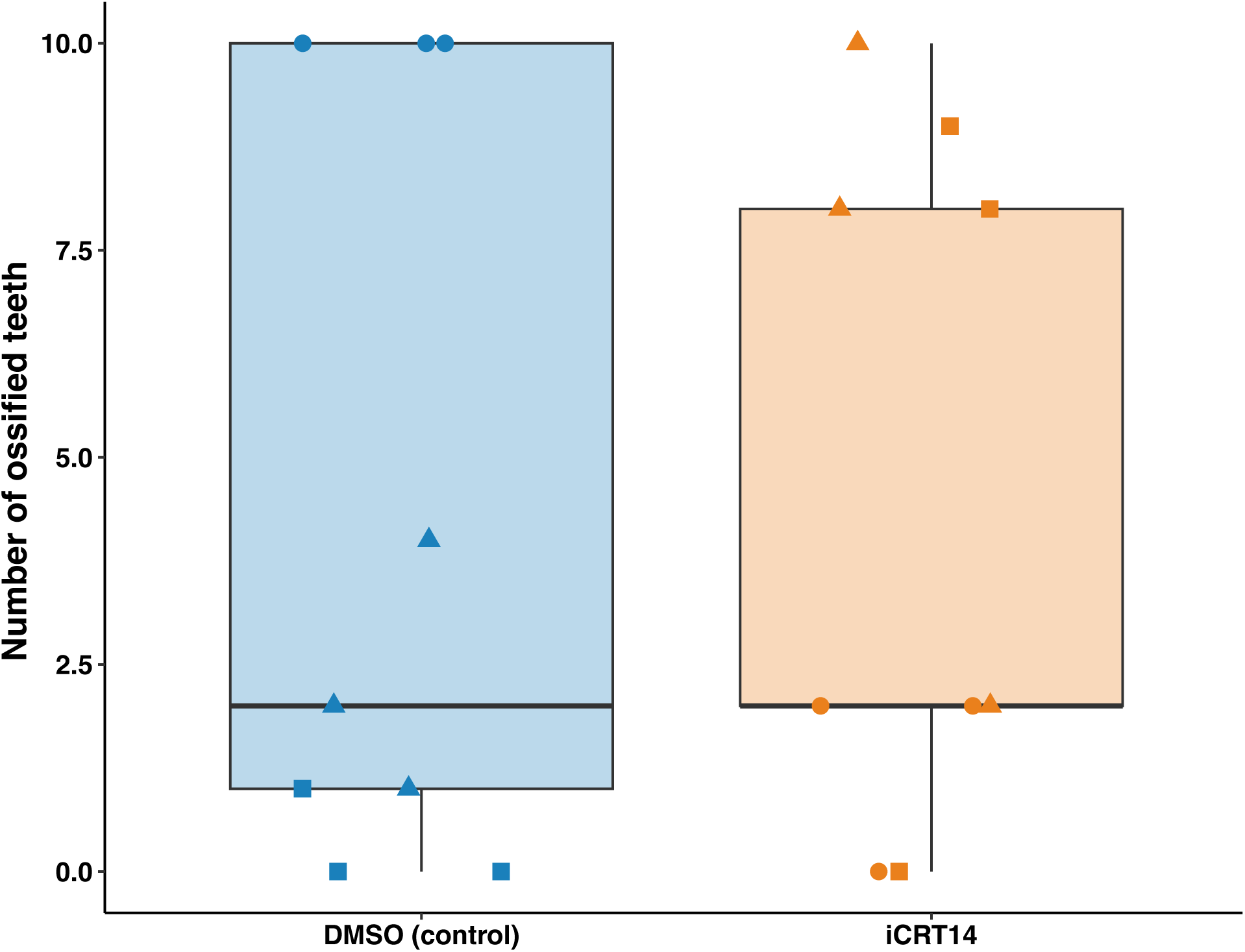
Number of ossified teeth was not affected by Wnt inhibition. Boxplot and jittered points of number of ossified teeth in *C. beltrani* larvae treated with either a control or Wnt inhibitor (iCRT14). Shapes represent different split-brood replicates.

